# Explainable protein-protein binding affinity prediction via fine-tuning protein language models

**DOI:** 10.64898/2026.03.30.715237

**Authors:** Harshit Singh, Rajeev Kumar Singh, Satya Pratik Srivastava, Suryavedha Pradhan, Rohan Gorantla

## Abstract

Protein–protein interactions underpin virtually every aspect of cellular life, and the precise quantification of their binding affinity is fundamental to understanding immune recognition, disease mechanisms, and the rational design of therapeutic antibodies. Yet predicting binding affinity at scale remains an unsolved challenge: reliable experimental assays are low-throughput and expensive, while computational methods that depend on three-dimensional complex structures cannot be applied to the vast majority of clinically relevant targets where structural data are absent. Here we present BALM-PPI, a framework that predicts protein–protein binding affinity from amino acid sequence alone. Both proteins are encoded by a protein language model trained on evolutionary sequence data and projected into a shared representational space, where their distance directly reflects binding strength. Fine-tuning this protein language model requires updating fewer than 1% of its parameters, and we show that this targeted adaptation steers the model toward interface-relevant sequence signals rather than spurious background correlations. On a curated benchmark of over 12,000 protein complexes, BALM-PPI matches or exceeds the accuracy of structure-based methods and retains predictive power for proteins with less than 30% sequence identity to the training set. Using only a subset of project-specific assay data, BALM-PPI outperforms a recent method trained on three times the data, suggesting that the model has already encoded the underlying interaction signals and requires only minimal supervision to specialise to a new target. BALM-PPI further provides residue-level attribution maps that pinpoint the amino acid positions driving each affinity prediction, consistently recovering experimentally validated interaction hotspots across enzyme-inhibitor, signalling, and antibody-antigen systems without any structural input during training. This allows predictions to be cross-validated against structural and mutagenesis evidence, providing a mechanistic basis for candidate shortlisting ahead of experimental follow-up. BALM-PPI is freely accessible via an interactive web server.

## 1 Introduction

Protein-protein interactions are the molecular basis of virtually every biological decision a cell makes, from mounting an immune response to propagating a growth signal or repairing damaged DNA. The strength with which two proteins associate, their binding affinity, determines whether an interaction is transient or stable, specific or promiscuous, and is therefore a central quantity in understanding disease mechanisms and in the design of therapeutic proteins such as monoclonal antibodies and biologics^1–3^.

Despite its importance, accurate affinity prediction at scale remains an unsolved problem. On the experimental side, established techniques such as surface plasmon resonance, isothermal titration calorimetry, and deep mutational scanning yield precise measurements but are inherently low-throughput and resource-intensive, making exhaustive variant screening impractical in early drug discovery. Computational methods offer a faster alternative, and structure-based approaches including Rosetta, FoldX, and molecular dynamics have long served as the gold standard when high-resolution complex structures are available^4–6^. Machine-learning extensions using graph neural networks and three-dimensional convolutional architectures, increasingly coupled with AlphaFold-multimer, have further improved accuracy^7^. Yet all structural methods share a fundamental ceiling: they require atomic-level complex structures as input, restricting throughput and precluding application to the vast majority of therapeutically relevant targets for which reliable structures remain unavailable.

Sequence-based approaches offer a scalable alternative, and the recent emergence of protein language models (PLMs) has substantially raised their ceiling. PLMs such as ESM-2, pre-trained on hundreds of millions of natural protein sequences, learn rich representations that implicitly encode structural, evolutionary and biochemical properties without ever observing a three-dimensional coordinate^8,9^. This makes them a natural foundation for sequence-only affinity prediction. Existing PLM-based methods typically concatenate embeddings from both proteins and pass the combined vector through a regression head^10^, a design that conflates interaction-specific signals with global sequence properties and limits generalisation across evolutionarily distant proteins and diverse assay technologies^11–13^. Methods such as MVSF-AB^14^ have made progress in antibody–antigen affinity prediction by combining multi-view sequence features, but still require large labelled datasets and do not learn a representation that transfers readily to new targets. Two other properties are essential for practical deployment in therapeutic programmes, yet are largely absent from current sequence-based methods. First, reliability under distribution shift, a model trained on one antigen class or protein family must retain predictive power when applied to evolutionarily distant scaffolds and novel target classes not seen during training, without catastrophic miscalibration. Second, explainability, residue-level rationales that can be cross-referenced against structural and mutagenesis data are essential for predictions to be acted upon with confidence in lead-optimisation campaigns, rather than treated as opaque scores that offer no mechanistic insight. A method that jointly addresses reliable affinity prediction, cross-family transferability, and explainability from sequence alone remains an open challenge.

Here we present BALM-PPI, a framework that addresses these gaps by recasting protein–protein binding affinity prediction as a metric learning problem. Rather than concatenating embeddings, BALM-PPI projects both interacting proteins into a shared latent space in which cosine distance directly encodes experimental binding affinity (Fig. 1). Parameter-efficient fine-tuning via Low-Rank Adaptation (LoRA)^15–17^ adapts the ESM-2 backbone while updating fewer than 1% of model parameters, steering the model toward interface-relevant sequence signals rather than spurious background correlations, while preserving the broad biological knowledge acquired during pre-training. Crucially, the same fine-tuning mechanism provides a natural route for adaptation to new targets: a small set of labelled measurements from any binding assay is sufficient to specialise the model to a new antigen without full retraining, enabling deployment across diverse experimental contexts with minimal additional data. BALM-PPI further accompanies each affinity prediction with residue-level attribution maps via Integrated Gradients^18^, pinpointing the amino acid positions that drive binding and consistently recovering experimentally validated interaction hotspots across enzyme–inhibitor, signalling, and antibody–antigen systems without structural supervision. This allows predictions to be cross-validated against structural and mutagenesis evidence, providing a mechanistic basis for candidate shortlisting ahead of experimental follow-up. The full prediction and explainability pipeline is made immediately accessible through an interactive web server (https://huggingface.co/spaces/Harshit494/BALM-PPI) and open-source code (https://github.com/rgorantla04/BALM-PPI.git).

**Figure 1.**
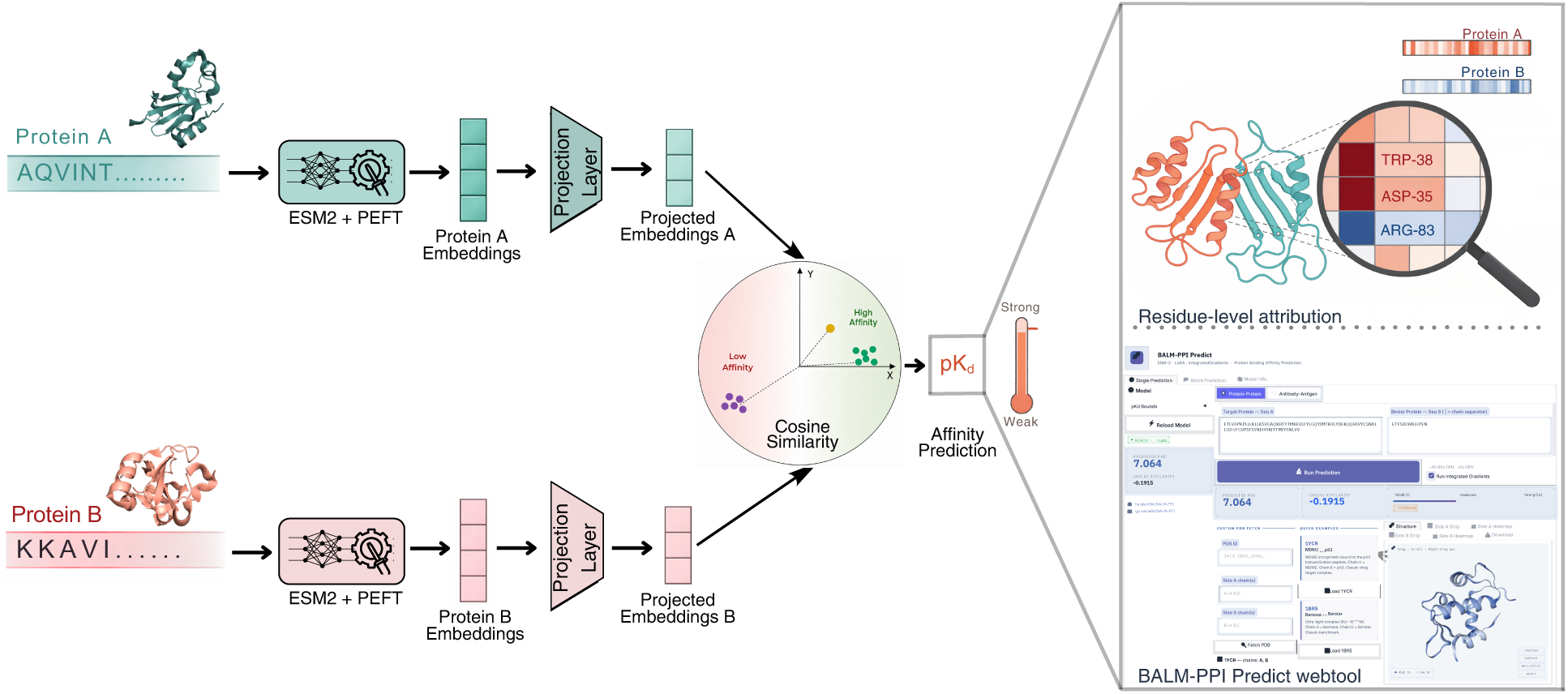
Overview of the BALM-PPI framework for explainable protein-protein binding affinity prediction. Protein A and Protein B sequences are independently processed using the ESM2 + PEFT encoder to generate contextual sequence embeddings. Parameter-efficient fine-tuning is achieved through LoRA adapters inserted into the frozen ESM-2 attention layers, enabling task-specific adaptation while keeping most pretrained parameters fixed. The generated embeddings are passed through projection layers to obtain representations in a shared latent space, where cosine similarity is used to estimate the binding affinity score (p*K*_*d*_). The framework additionally provides residue-level explainability using Integrated Gradients (IG), highlighting important interface residues through heatmaps and 3D structural visualizations. A web-based interface allows users to input raw protein sequences and obtain predicted affinity scores, cosine similarity values, interactive structural visualizations, attribution maps, and downloadable analysis outputs including IG score CSV and summary JSON files.

## 2 Results

### 2.1 BALM-PPI generalises to evolutionarily distant proteins under low sequence similarity

We trained and evaluated BALM-PPI on the PPB-Affinity dataset^19^, the largest publicly available collection of curated protein-protein binding affinity measurements, comprising 12,019 complexes with experimental *K*_*d*_ values spanning enzyme-inhibitor, signalling, TCR-pMHC and antibody-antigen interactions drawn from SKEMPI v2.0, SAbDab, PDBbind v2020, ATLAS and the Affinity Benchmark v5.5. Dissociation constants were converted to *pK*_*d*_ = − log_10_(*K*_*d*_) to obtain a uniform target distribution, and all results are reported as mean and standard deviation over five-fold cross-validation, using RMSE for absolute error and Pearson and Spearman correlations for relative agreement with experiment.

To probe how the model behaves as the gap between training and test proteins widens, we evaluated three splits of increasing difficulty. The random split partitions complexes uniformly and measures interpolation within the training distribution, but is known to over-estimate generalisation when redundant complexes appear in both folds^20^. The cold split groups by parent PDB so that no complex appears in both training and test, and measures generalisation to unseen interfaces. The sequence-similarity split applies agglomerative clustering on *K*-mer Jaccard distance and enforces a strict 30% identity ceiling between training and test proteins, the regime in which most therapeutically relevant targets sit and in which sequence-only predictors typically collapse. We compared three model configurations on every split: a concatenation regressor that joins both ESM-2 embeddings and passes them through an MLP head, a cosine-distance model on the same backbone without parameter-efficient adaptation, and the full BALM-PPI with LoRA adapters on the attention layers. The first two variants isolate the contribution of the metric-learning geometry and the third quantifies the additional gain from adapting the encoder.

On the random split, BALM-PPI reaches Pearson 0.887 ± 0.008, Spearman 0.886 ± 0.008 and RMSE 0.994±0.025 p*K*_*d*_ units, compared with RMSE 1.345±0.029 for the concatenation regressor and 1.192 ± 0.022 for the cosine model without LoRA (Fig. 2a, Supplementary Table S2). On the cold split, where information leakage from shared complexes is removed, BALM-PPI reaches Pearson 0.730 ± 0.025, Spearman 0.715 ± 0.046 and RMSE 1.486 ± 0.115, against 1.631 ± 0.022 for the regressor and 1.545 ± 0.068 for the cosine model without LoRA. On the strictest sequence-similarity split, BALM-PPI reaches Pearson 0.612 ± 0.086 at RMSE 1.672 ± 0.195, against Pearson 0.555 ± 0.097 at RMSE 1.755 ± 0.160 for the regressor. Performance therefore decays gradually rather than catastrophically as the distribution shift intensifies, and rank correlation degrades more slowly than absolute error, which indicates that the latent space preserves relative binding preferences even when the affinity scale becomes harder to calibrate.

**Figure 2.**
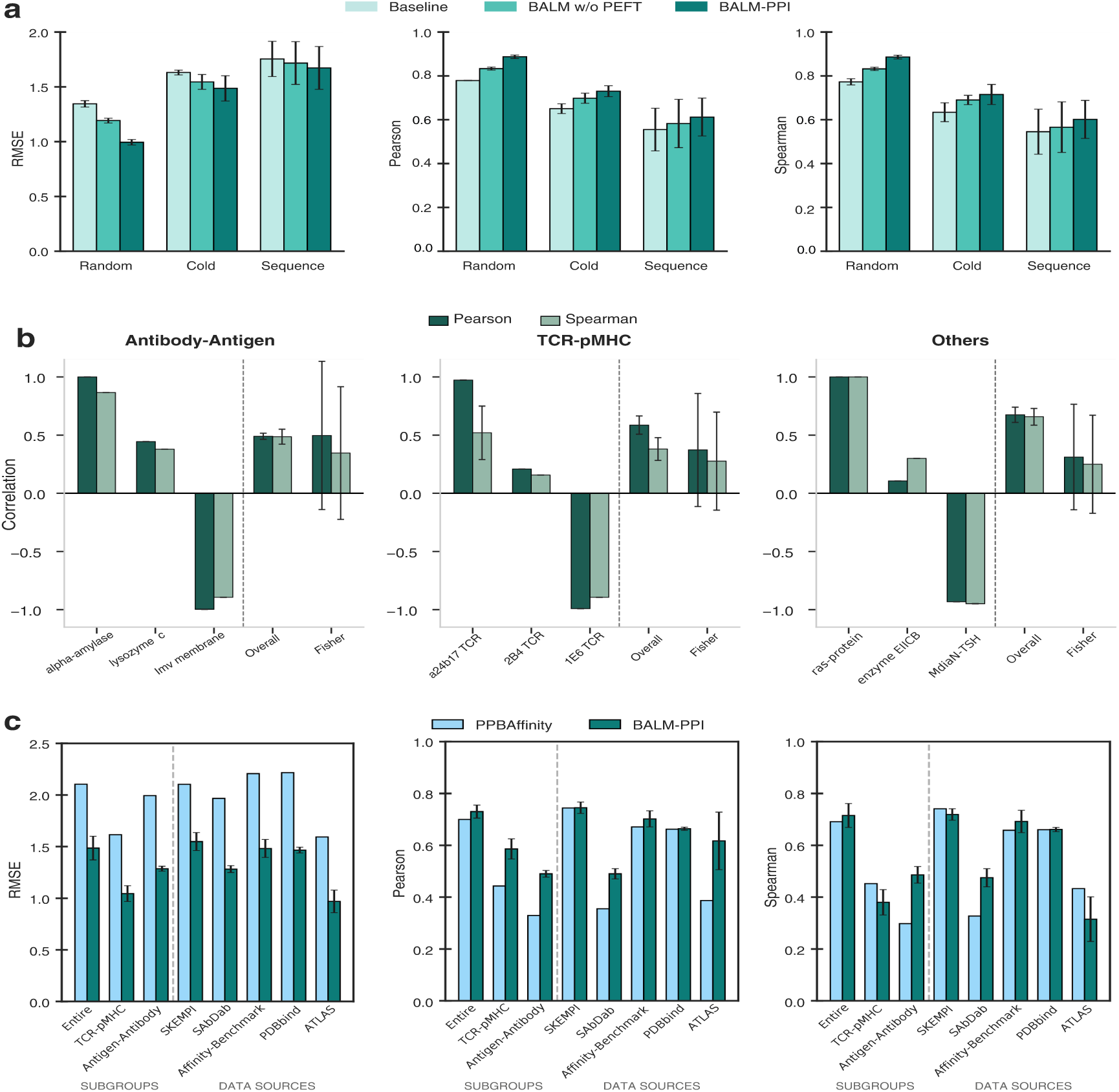
Performance of BALM-PPI under increasing distribution shift and comparison with existing baselines. **a)** Aggregate predictive performance of BALM-PPI under three cross-validation settings: random split, cold split, and sequence-similarity split. Bars report root mean squared error (RMSE; lower is better) and Pearson and Spearman correlations (higher is better) for the Baseline model, BALM without PEFT, and BALM-PPI, illustrating increasing difficulty under distribution shift and the incremental gains from each modeling component. Error bars show mean ± s.d. across 5 folds **b)** Biological deep-dive into generalization for antibody–antigen, TCR–pMHC, and general protein complexes (Others). For each subgroup, three representative individual receptors illustrate the model’s performance spectrum (top-performing, median, and low-performing targets). Aggregate performance is reported as Overall (mean correlation across all pairs in the subgroup) and Fisher (Fisher Z-transformed average of target-specific correlations). **c)** Comparison of sequence-only BALM-PPI with the structure-based PPB-Affinity baseline across biological subgroups (left of dashed line) and dataset sources (right of dashed line). Performance is reported using RMSE, Pearson, and Spearman metrics for the full dataset, immune-related subsets, and individual benchmark sources. BALM-PPI consistently reduces error and achieves stronger correlations across most subsets, highlighting robustness to dataset shift and competitive accuracy relative to structure-based methods.

To validate our choice of backbone encoder, we conducted an ablation study comparing ESM-2 against ProGen-2 (small and medium)^21^, AbLang-2^22^ and ESM-C^23^ under the cold split regime for BALM-PPI without PEFT, each paired with projection heads of 256, 512 and 1024 dimensions. ESM-2 delivered consistent performance across all projection sizes as demonstrated in (Supplementary Fig. S3) and (Supplementary Table. S7), confirming that its representations are best suited for addition-space projection. This motivates its use as the default backbone throughout the main experiments. Two patterns are consistent across all three splits. The first is the ordering of the three configurations. Replacing concatenation with the cosine-distance objective is responsible for the larger share of the gain on every split, and adding LoRA contributes a further consistent improvement on top of it by updating fewer than 1% parameters of the backbone weights. The second is that standard deviations are small under the random split and grow under the cold and sequence-similarity splits, indicating stable optimisation in-distribution and predictable variability under out-of-distribution evaluation.

Aggregate metrics conceal where prediction is reliable and where it is not. Under the cold split, target-specific correlations (Fig. 2) are high for many unseen targets in the antibody-antigen, TCR-pMHC and general protein subgroups, but a minority of difficult targets show weak or inverted ranking. We therefore report both the pooled correlation and the Fisher *Z*-averaged target-level correlation, together with the per-target spread shown by the error bars in Fig. 2b. The two statistics can disagree, and reporting them jointly captures the full spectrum from successful generalisation to reverse-ranking failure, which we consider a more rigorous summary of out-of-distribution behaviour than either statistic in isolation. The negative-correlation cases are concentrated on scaffolds that are evolutionarily distant from anything in the training set and indicate that, when transfer fails, it fails by mis-scaling the affinity ranking rather than by missing the interface signal. The two sections that follow show that this failure mode is correctable with a small amount of in-domain data.

### 2.2 BALM-PPI surpasses structure-based deep learning baseline across biological subgroups on PPB-Affinity

We benchmark BALM-PPI against the PPB-Affinity structure-based deep learning baseline under the cold split regime to quantify how far a PLM-driven cosine similarity objective can go without access to explicit 3D complex structures. In addition to overall performance on the full evaluation set, we compare our model’s results for two subgroups (TCR-pMHC and antibody-antigen) and provide a source-wise breakdown across the constituent PPB-Affinity data sources to assess cross-domain robustness as shown with comparison bars in Fig. 2 (Detailed tables with results and comparison is provided in Supplementary Tables S3 and S4). On the full evaluation set, *BALM-PPI* reduces prediction error to an RMSE of 1.486 ± 0.115, compared with 2.104 for the PPB-Affinity baseline, while maintaining strong Pearson and Spearman correlations, suggesting that the model is learning biologically meaningful interaction signals.

For biological subgroups, substantial gains were observed in both antibody-antigen and TCR-pMHC complexes, indicating that the framework effectively identifies interface-relevant signals from sequence embeddings. This performance remained consistent across diverse data repositories, with *BALM-PPI* achieving lower RMSE and higher Pearson correlation across SKEMPI v2.0, SAbDab, PDBbind v2020, ATLAS, and the Affinity Benchmark v5.5. These findings suggest that the BALM-PPI framework offers strong predictive power by learning biologically meaningful signals across a range of data sources and subgroups without requiring explicit 3D structural input.

### 2.3 Few-shot adaptation on small subset of single-mutational AB-Bind data surpasses models trained on 90% data

To assess out-of-distribution generalisation and adaptation efficiency, we evaluated our framework on the independent AB-Bind dataset^24^, following the benchmarks established by MVSF-AB^14^. We enforced a strict zero-shot environment by removing any AB-Bind complexes that overlapped the PPB-Affinity training set at the level of parent PDB identifier or identical antigen–antibody sequence pair, and reduced the number of complexes from 1,089 to 272 non-redundant complexes.

Zero-shot evaluation on this cleaned set yields RMSE of 1.217 but negative correlation, due to scale mismatch between PPB-Affinity-trained affinity range and the AB-Bind mutational landscape, This is an expected result due to strict de-overlapping that enforces a truly out-of-distribution test set with single point mutation.

Few-shot adaptation rapidly achieves strong positive correlation as shown in (Table 1): With a small set of AB-Bind data, BALM-PPI yields Pearson *r* = 0.613 ± 0.120, and with only 30% of data, it yields *r* = 0.756 ± 0.040 with RMSE = 0.688 ± 0.035. This surpasses MVSF-AB^14^ (Pearson of 0.739, RMSE of 1.905), which was trained on 90% of the complete, non-de-overlapped dataset, highlighting a significant improvement in data efficiency.

**Table 1.** AB-Bind performance on the cleaned dataset: zero-shot baseline and few-shot adaptation results (mean ± s.d. over 3 random seeds: 42, 123, 999).

| Training Regime | Data Fraction | RMSE ↓ | Pearson ↑ | Spearman ↑ |
| --- | --- | --- | --- | --- |
| Zero-shot | 0% | 1.217 | -0.337 | -0.349 |
| Few-shot | 10% | $0.859 \pm 0.115$ | $0.613 \pm 0.120$ | $0.570 \pm 0.103$ |
| Few-shot | 20% | $0.750 \pm 0.052$ | $0.727 \pm 0.062$ | $0.682 \pm 0.062$ |
| Few-shot | 30% | **$0.688 \pm 0.035$** | **$0.756 \pm 0.040$** | **$0.712 \pm 0.052$** |

### 2.4 Cross-domain generalization of BALM-PPI on diverse AbBiBench assays

Using nine deep-mutational-scanning (DMS) antibody–antigen assays from the AbBiBench dataset^25^, we assessed cross-domain performance, as detailed in Supplementary Table S5. Each assay corresponds to a distinct antibody–antigen system with its own parental sequence, mutational distribution, and label scale. This provides a stringent test of whether the learned latent space captures fine-grained mutational effects beyond coarse interaction propensity.

Zero-shot performance varies across assays, reflecting compound distribution shift in antigen class, antibody scaffold and experimental label scale. Few-shot adaptation with 10–30% of each assay consistently improves correlation, with the largest gains on influenza hemagglutinin stem-binding assays (e.g., CR6261, CR9114; Pearson rising from ∼0.3 to >0.95) and smaller but consistent gains on SARS-CoV-2 HR2 assays, suggesting harder domain shift or lower signal-to-noise in the latter. Assay-level performance across zero-shot and few-shot settings with different metrics is demonstrated in Fig. 3, while Supplementary Table S6 reports the full RMSE, Pearson and Spearman results for all nine assays and data fractions, and Fig. S2 in Supplementary information shows the predicted-versus-experimental scatter plots across training fractions.

**Figure 3.**
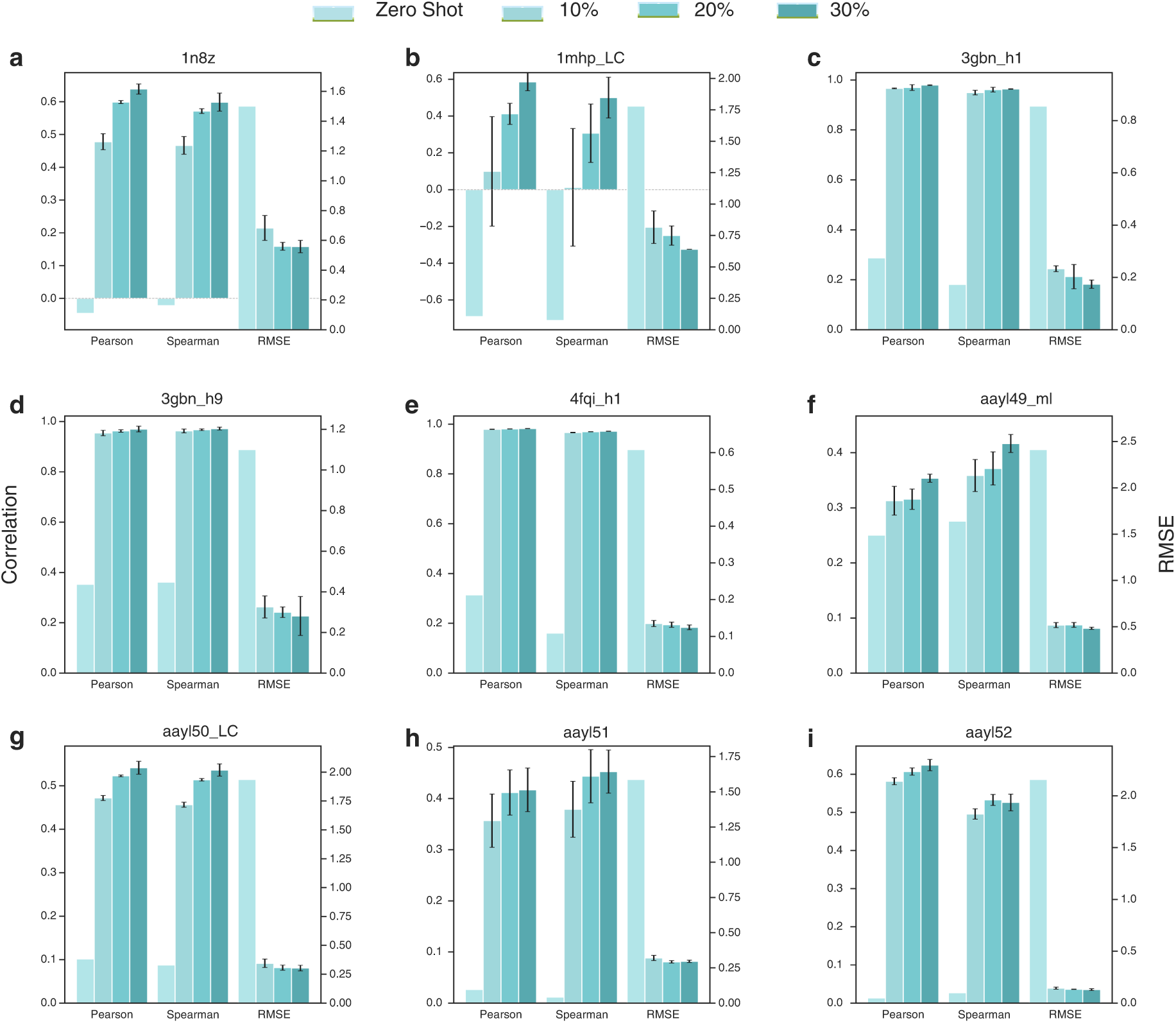
Generalization and few-shot adaptation on nine deep-mutational-scanning AbBiBench assays. (a-i) Zero-shot and few-shot evaluation on 9 representative AbBiBench deep-mutational-scanning assays. For each assay, the bar plots summarize Pearson, Spearman, and RMSE across zero-shot and few-shot adaptation settings (10%, 20%, 30%). Bars show mean ± s.d. of 3 random seeds (42, 123, 999). Consistent performance gains with limited assay data demonstrate rapid calibration of the shared latent representation to new antibody–antigen mutational landscapes without full fine-tuning.

Notably, assays with near-zero or negative zero-shot correlation become strongly positive after few-shot learning, indicating that model can adapt quickly to various downstream tasks. Performance gains from 10% to 30% data exhibit diminishing returns, consistent with a representation-learning regime in which most biophysical priors are already encoded and only minimal supervision is required to specialise to new mutational neighbourhoods. From a practical perspective, experimental campaigns could leverage our framework for rapid prioritisation after profiling only a small variant subset.

Few-shot learning yields substantial improvements with only 10-30% of labeled variants, consistent with the interpretation that most task-specific differences arise from assay-dependent scaling and noise rather than a need to relearn interaction features from scratch.

The consistent directionality of improvement across heterogeneous antigen classes supports the hypothesis that BALM-PPI learns a transferable interaction geometry that requires only scale alignment to new experimental setup.

Assays targeting influenza hemagglutinin stem epitopes exhibit strong few-shot gains, consistent with partially shared structural or sequence-level motifs captured during pretraining. In contrast, assays involving SARS-CoV-2 display more modest improvements, potentially reflecting distinct epitope geometry, narrower dynamic range, or increased experimental noise. These differences suggest that zero-shot transfer performance is influenced not only by evolutionary distance but also by assay-specific signal-to-noise characteristics and mutational landscape smoothness. The observation that near-zero or negative zero-shot correlations can become strongly positive after limited adaptation suggests that the primary bottleneck is calibration mismatch rather than absence of transferable interaction features. In other words, the latent representation appears to encode relevant mutational effects, but their mapping to assay-specific binding scales requires minor reparameterization. This interpretation aligns with the effectiveness of PEFT-only fine-tuning, which adjusts low-rank adapter weights without modifying the core protein language model. Performance improvements from 10% to 30% training data exhibit diminishing returns, indicating that a relatively small fraction of labeled variants suffices to align the shared embedding space to new antibody-antigen systems. From a practical perspective, this suggests that experimental campaigns could leverage BALM-PPI for rapid prioritization after profiling only a small subset of variants. Overall, the combined evidence from in-distribution evaluation, robustness tests, and few-shot transfer indicates that BALM-PPI offers a strong accuracy and data-efficiency trade-off, and delivers competitive performance without requiring structural inputs, and can be specialized quickly to new antibody-antigen systems with minimal additional supervision.

### 2.5 Residue-level attribution recovers interaction hotspots from sequence

To explain the learned representations and identify amino acid residues contributing most to the predicted protein-protein binding affinity, we performed a comprehensive explainability analysis of the fine-tuned *BALM-PPI* model. The analysis was designed to assess whether the model’s predictions align with biologically meaningful residue-level interactions.We performed attribution analysis on five protein-protein interactions and four antibody-antigen complexes to evaluate the biological importance of our model’s predictions.

To evaluate whether BALM-PPI identifies right interactions from sequences while learning, we analyzed the Barnase-Barstar complex (PDB: 1BRS) ^26^. This system is a well-characterized model for electrostatic optimization, exhibiting a near-diffusion-limited association rate driven by charge-charge complementarity at the interaction interface ^27^. In the zero-shot state, the model’s attribution heatmaps (Fig. 4) highlight several residues known to be energetic hotspots for this interaction. This indicates that ESM-2 representations, when projected through the cosine similarity framework, can capture underlying electrostatic features relevant to protein docking without task-specific fine-tuning. The attribution analysis (predicted *pK*_*d*_ = 8.79) shows that *BALM-PPI* concentrates importance on a compact set of residues on the Barstar (inhibitor) chain (deepsalmon): *Trp-38, Asp-35, Asn-33, Glu-76, and Thr-42*. The identification of acidic residues (*Asp-35* and *Glu-76*) along with polar stabilizers (*Asn-33* and *Thr-42*) and the aromatic residue *Trp-38* aligns with established ground-truth data showing that Barstar is optimized for electrostatic binding to Barnase ^27^. On the Barnase (enzyme) chain (cyan), the highest importance scores are assigned to *Arg-59, Arg-83, Arg-87, and His-102*. These positively charged side chains complement the negatively charged residues identified on Barstar. This pattern is consistent with the known binding mechanism. The alignment between these residue-level attributions and the known biophysical properties of the Barnase-Barstar interface suggests that the zero-shot model identifies physically relevant interaction determinants rather than relying solely on global sequence correlations.

**Figure 4.**
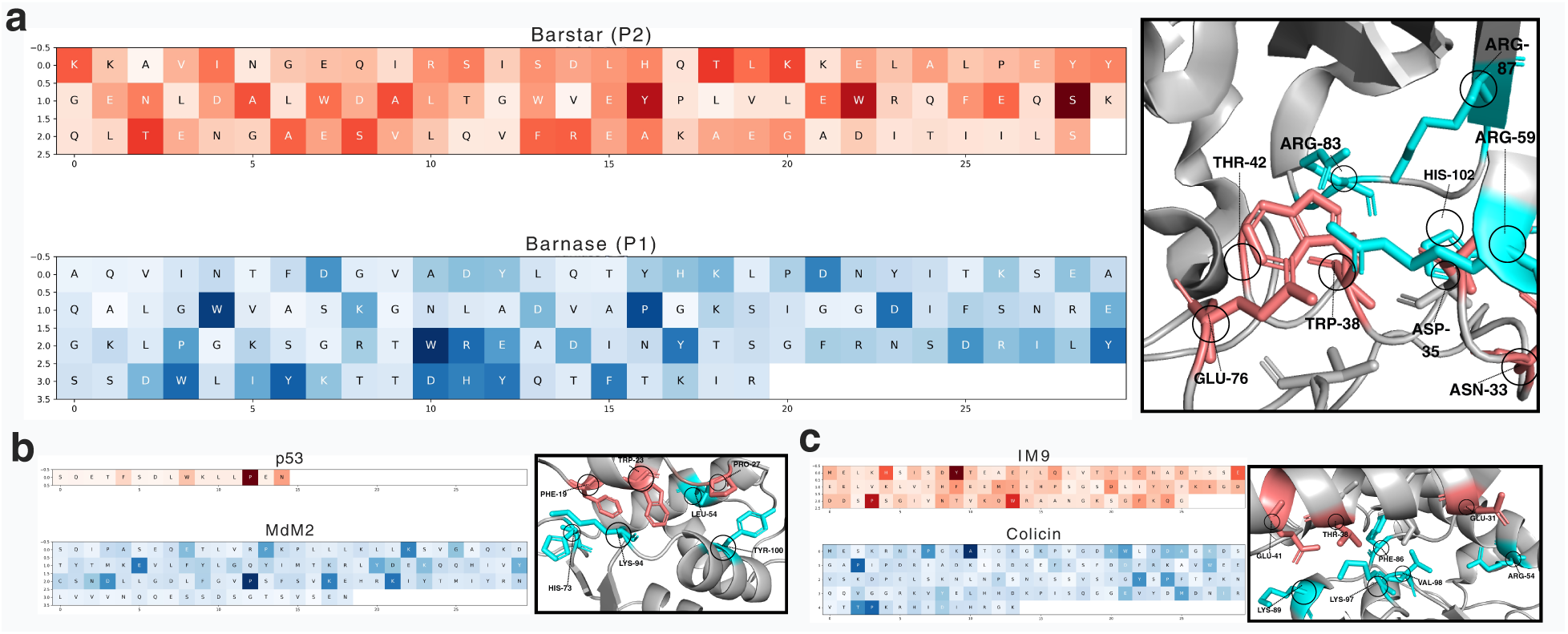
Zero-shot residue-level attribution maps for diverse protein-protein interaction mechanisms. Integrated Gradients (IG) attributions from the zero-shot BALM-PPI model are mapped onto five representative complexes to visualize which residues most strongly influence the predicted binding affinity. For each subfigure, the two interaction partners are shown in **cyan** and **deepsalmon**. The panels cover distinct recognition regimes: (a) Barnase-Barstar (electrostatically optimized enzyme-inhibitor binding), (b) MDM2-p53 (peptide-in-groove recognition) (c) Colicin-IM9 (ultra-high-affinity toxin-immunity binding). The consistent localization of attribution to known interface regions indicates that the sequence-only model recovers physically meaningful interaction hotspots without explicit 3D supervision.

For the Ras-Raf complex, BALM-PPI predicts a *pK*_*d*_of 9.06. We analyzed this interaction to evaluate the model on complex regulatory interfaces characterized by an extended binding surface ^28^. In the zero-shot attribution analysis (Fig. S5), BALM-PPI highlights a continuous binding surface that aligns with the structural interface described in literature ^28^. On the Ras (GTPase) chain (cyan), the highest importance scores are assigned to *Val-21, Glu-31, Asp-33, Ile-36, Glu-37, Tyr-40, and Glu-76*. These residues are located within the canonical interaction surface and define the primary binding patch. On the Raf chain (deepsalmon), attribution is concentrated on *Pro-63, Lys-65, Arg-67, Val-69, Val-70, Asn-71, Lys-84, Lys-87, and Val-88*. This residue-level pattern aligns with energetic analyses and per-residue free-energy decompositions performed on Ras-Raf complexes ^29^. These results suggest that BALM-PPI identifies residues and side-chain chemistries associated with the stabilization of the complex.

We evaluated the Trypsin-BPTI complex (PDB: 3BTK), a system characterized by a high degree of steric fit and rigid binding conformations maintained by disulfide-stabilized loops. For this complex, BALM-PPI predicts a *pK*_*d*_ of 8.32. The zero-shot attribution analysis (Fig. S5 shows that the model highlights residues associated with the structural rim and the environment supporting the binding interface, rather than focusing solely on catalytic sites. The attribution heatmap for the Trypsin chain (cyan) shows a focus on residues that define the canonical binding pocket (Fig. S5). The model assigns high importance to *Asp-189* and *Gln-192*, which are located in the substrate-binding pocket and are known to influence binding specificity and strength. Additionally, the model highlights *Tyr-39* and *His-40* at the interface, as well as *Lys-515*, a residue associated with longer-range electrostatic interactions that contribute to complex stability. On the BPTI chain (deepsalmon), the model focuses on *Pro-513* and the basic residues *Arg-517* and *Arg-539*. This pattern aligns with structural analyses indicating that these residues contribute to stabilization through shape complementarity and electrostatic effects ^30^. These results show that the attribution scores correspond to residues previously identified as important for the energetic profile of the Trypsin-BPTI interaction.

To evaluate the model on different molecular geometries, we analyzed the MDM2-p53 regulatory complex (PDB: 1YCR), for which BALM-PPI predicts a *pK*_*d*_of 8.64. This interaction involves a segment of the p53 transactivation peptide that binds within a hydrophobic pocket on the MDM2 receptor.^31^ The zero-shot attribution analysis (Fig. 4) shows that BALM-PPI highlights the p53 hotspot residues *Phe-19* and *Trp-23*, along with *Pro-27*. This attribution pattern is consistent with reported structure-affinity relationships for MDM2 recognition ^32^. On the MDM2 receptor side (cyan), the model highlights *His-73, Lys-94, Leu-54, and Tyr-100*. The identification of *Leu-54* and *Tyr-100* is consistent with their roles in forming the walls of the binding cleft. On the p53 peptide side (deepsalmon), the highest importance scores are assigned to *Phe-19* and *Trp-23*, which function as hydrophobic anchors. *Pro-27* is also identified, which is a residue known to influence peptide helicity and docking geometry ^32^. The alignment between the attribution heatmap (Fig. 4) and known structural determinants suggests that BALM-PPI identifies residues relevant to hydrophobic peptide-protein recognition in a zero-shot setting.

For the Colicin-IM9 toxin-antitoxin complex, BALM-PPI predicts a *pK*_*d*_ of 8.83. We analyzed this system to evaluate the model on an interaction characterized by extremely high binding affinity (*pK*_*d*_ ≈ 16) and high specificity between cognate partners ^33,34^. The zero-shot attribution analysis (Fig. 4) shows that the model highlights a set of residues corresponding to the electrostatic and shape-complementary features of the binding interface. On the Immunity Protein (IM9) (deepsalmon), BALM-PPI identifies residues within a known acidic patch, specifically *Glu-31, Thr-38, and Glu-41*. On the Colicin side (cyan), the highest importance scores are assigned to *Arg-89, Lys-89, Phe-89, Lys-97, and Val-98*. This set combines basic side chains (Arg/Lys) with hydrophobic residues (Phe/Val), which is consistent with a binding mode where charge complementarity is reinforced by local packing in the interfacial region. The alignment between these highlighted residues and established structural determinants ^34^ indicates that BALM-PPI identifies relevant interfacial features for this high-affinity complex in a zero-shot setting.

For Antibody-Antigen, We analyzed the interaction between the antibody CR6261 and the Influenza A Hemagglutinin (HA) stem (PDB: 3GBN) ^35^. CR6261 neutralizes several influenza subtypes by binding to a conserved hydrophobic pocket in the HA stem, primarily using its Heavy chain CDRH2 and CDRH3 loops. We compared the attribution maps in two states: Zero-shot (pretrained on the PPB-Affinity dataset) and few-shot (fine-tuned on 30% of the CR6261-H1HA assay data). As shown in Fig. 5, the zero-shot model identifies the general area of the binding interface, which suggests that the backbone representations contain information relevant to protein contact surfaces. Following few-shot fine-tuning, the attribution scores for specific residues at the hydrophobic interface increase, which narrows the identified hotspots to residues associated with the interaction. The attribution hotspots align with the canonical HA-stem interface bound by CR6261 ^35^ (Fig. 5). On the CR6261 antibody side, the highest importance scores are assigned to *Phe-29, Arg-30, Tyr-32, Ala-33, Asp-73, and Tyr-98*. These residues are involved in hydrophobic packing and stabilization within the heavy-chain binding region. On the H1N1 HA side, the model highlights *Val-40, Leu-42, Gln-42, Val-52, His-18, His-38, Asp-19, Asp-46, Ser-291, and Phe-294*. These residues correspond to the conserved stem epitope and its immediate environment. The predicted *pK*_*d*_ for this complex was 8.40 in the zero-shot setting and 9.37 after few-shot adaptation. Across the antibody cases, a consistent pattern is observed: zero-shot attributions identify the general interface region, while few-shot adaptation increases the attribution contrast at specific interface residues.

**Figure 5.**
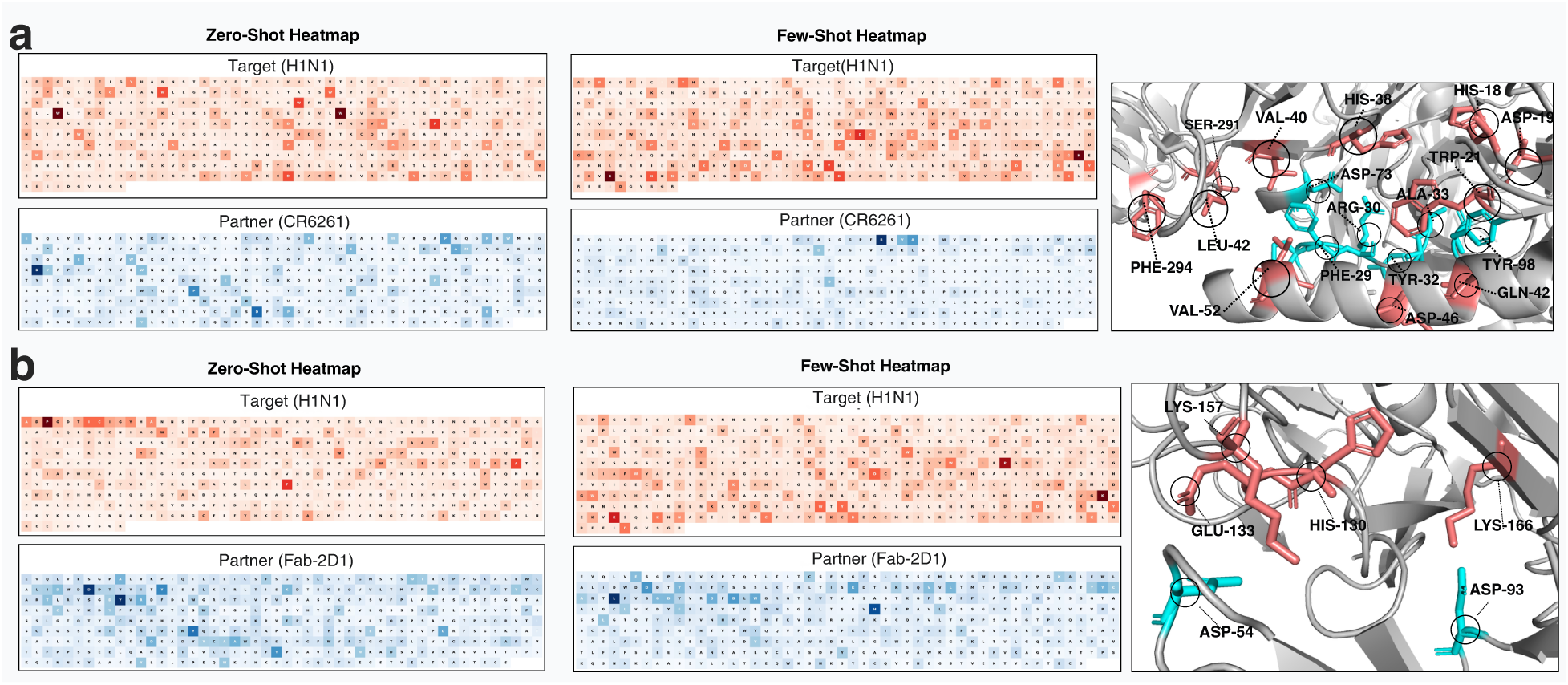
Residue-level attribution maps for antibody-antigen recognition. IG attributions from BALM-PPI are mapped onto representative antibody-antigen complexes to visualize residues that most strongly drive the predicted affinity signal. In each panel, the antibody is shown in **cyan** and the antigen in **deepsalmon**. The panels cover influenza hemagglutinin stem- and head-targeting antibodies and a canonical protein antigen system: (a) CR6261-H1N1 (PDB: 3GBN), (b) Fab-2D1-H1N1 (PDB: 3LZF). In the zero-shot setting, attribution localizes to contiguous paratope/epitope regions; few-shot PEFT further sharpens these hotspots for assay-specific landscapes.

We further evaluated the model on the interaction between the neutralizing antibody CR9114 and H1N1 hemagglutinin ^36^. CR9114 binds a conserved stem epitope and is characterized by specific structural requirements for pan-influenza protection ^37^. In the CR9114 heatmap (Fig. S6), the highest antibody-side attributions are assigned to *Phe-54, Ile-34, Asn-31, Pro52A, Tyr-98, Tyr-100, and Tyr-100A*, along with a cluster including *Ala-71, Asp-72, Ile-73, and Phe-74*. This pattern is consistent with the reported binding mode where a hydrophobic insertion residue (*Phe-54*) is supported by an aromatic cap (*Tyr-98/Tyr-100/Tyr-100A*) and scaffold residues that stabilize the paratope geometry. The correspondence between these attribution hotspots and structural analyses of CR9114 indicates that BALM-PPI identifies residues associated with broad protection from sequence data ^37^. The predicted *pK*_*d*_ for this complex was 8.87 in the zero-shot setting and 8.93 after few-shot adaptation.

We evaluated the interaction between Fab-2D1 and the 1918 H1N1 hemagglutinin head domain ^38^. This interaction differs from the hydrophobic targeting seen in stem-binding antibodies, as 2D1 binds to an electrostatic interface on the HA head domain ^38^. Comparison of the zero-shot and few-shot attribution maps (Fig. 5) shows a shift in the regions identified by the model. In the zero-shot state, the model identifies the heavy chain as the primary interactor. After few-shot adaptation, attribution scores increase for residues within the antigen’s 130-loop and 150-loop, which are the regions where the 2D1 antibody contacts the antigen ^38^. The residues identified by BALM-PPI align with known determinants of Fab-2D1 recognition (Fig. 5). On the Fab-2D1 side, the highest importance scores are assigned to *Asp-54* and *Asp-93*. The identification of these acidic residues is consistent with an interaction mode where electrostatics and salt-bridge formation contribute to stabilizing the complex. On the H1N1 HA side, the model highlights *His-130, Glu-133, Lys-157, and Lys-166*. The attribution for *Lys-157*, *Glu-133*, and *Lys-166* corresponds to residues involved in charge complementarity, while *His-130* is a residue associated with the local environment at the epitope. The correspondence between these attribution hotspots and published mutation analyses related to viral escape indicates that BALM-PPI identifies epitope and paratope residues from sequence data ^39^. The predicted *pK*_*d*_ for this complex was 8.48 in the zero-shot setting and 8.49 after few-shot adaptation.

We analyzed the HyHEL-10 antibody in complex with hen egg-white lysozyme to evaluate the framework on a non-viral protein scaffold. This system is characterized by aromatic packing and solvent exclusion at the interface, which are known to influence binding energetics ^40^. The attribution heatmap (Fig. S6) shows that higher importance scores are localized to the paratope-epitope contact region rather than distributed across the full protein surfaces. On the HyHEL-10 antibody side, the highest attributions include *Asp-32* and a cluster of aromatic and polar residues: *Asn-31, Tyr-33, Tyr-50, Ala-51, Gln-53, Tyr-53, Tyr-58, Tyr-96, Lys-97, and Trp-98*. The identification of multiple tyrosines and a tryptophan aligns with the role of aromatic packing and water exclusion in this interaction ^40^. Additionally, the attribution to *Asp-32* and *Lys-97* corresponds to residues involved in electrostatic anchoring and charge complementarity. On the Lysozyme side, the model highlights *His-15, Arg-21, Thr-89, Asn-93, Lys-96, Ile-98, Ser-100, Asp-101, and Gly-102*. These residues define the epitope patch, with *Arg-21* and *Lys-96* providing positive charges that complement the antibody’s *Asp-32*, while *Asp-101* is associated with the polar and electrostatic structure of the interface. The alignment between the residues identified in the heatmap (Fig.S6) and established hotspot analyses based on water-exclusion models indicates that BALM-PPI identifies residues relevant to the binding determinants of this antibody-antigen system ^40^. The predicted *pK*_*d*_for this complex was 8.77 in the zero-shot setting and 8.84 after few-shot adaptation.

The attribution analysis across five general protein-protein interactions and four antibody-antigen systems shows that the highest importance scores are consistently localized to interface regions. These identified residues align with previously reported mechanistic determinants of binding. The application of PEFT increases the attribution contrast at specific residues, which is consistent with the refinement of representations for assay-specific mutational landscapes. The identified hotspots correspond to various physical drivers of interaction, such as electrostatic complementarity in the Barnase-Barstar system, hydrophobic anchoring in the MDM2-p53 complex, and pocket-mediated recognition in Trypsin-BPTI. The model highlights residues with known functional roles, including conformational determinants like Pro-27 in p53 and residues within binding loops like His-130 in Fab-2D1. These results suggest that the latent-space similarity learning in the BALM-PPI framework identifies residue-level signals consistent with structural data, which may assist in identifying interaction hotspots for protein and antibody engineering.

### 2.6 Interactive web server for sequence-based affinity prediction and explainability

To make BALM-PPI immediately accessible without requiring local computational infrastructure, we developed an interactive web server, BALM-PPI·predict, that demonstrates the prediction and explainability pipeline through a browser interface. Users can provide raw amino acid sequences or fetch the FASTA sequences using PDB ID and chain IDs for the target and binder proteins.

Model weights are loaded from the public Hugging Face checkpoint by default, user can fetch the sequences using PDB IDs, and chain IDs. Upon submitting a prediction, the tool returns the predicted affinity value, cosine similarity score, and a binding strength gauge calibrated to the dataset’s *pK*_*d*_range. Integrated Gradients attributions are computed by enabling Run Integrated Gradients checkbox and visualised through a 3D NGL Viewer with residues coloured by attribution intensity, a residue strip heatmap showing per-residue IG scores, and a ranked bar chart of the top contributing residues for each protein. All results are available for download as an IG scores CSV and a summary JSON file. The web interface is illustrated in Fig.S4.Users can run batch predictions too by using the given format in CSV for both protein-protein and antigen-antibody.

## 3 Discussion and concluding thoughts

BALM-PPI shows that protein-protein binding affinity can be predicted from sequence alone at a level that matches structure-based methods on a benchmark of more than 12,000 complexes, retains predictive power for evolutionarily distant proteins under low sequence similarity to the training set, and recovers experimentally validated interaction hotspots without structural supervision. The combination matters because the targets where computational prediction is most needed are the same ones where reliable complex structures do not exist, and where current structure-based predictors therefore do not apply. For therapeutic antibody optimisation in particular, where experimental measurements are expensive and sparse and where structural coverage of the antigen of interest is the exception rather than the rule, a sequence-only predictor that comes with per-residue rationale is the kind of tool that can be inserted into a lead-optimisation campaign rather than treated as an external benchmark. The geometry is what makes the method work. Casting affinity as a cosine distance in a shared latent space pushes the disentanglement of interaction signal from bulk sequence statistics into the embedding step, where the pre-trained ESM-2 backbone has the right inductive bias, rather than leaving it to a regression head sitting on top of concatenated embeddings. Across the architectural variants we evaluated, the cosine objective alone closes most of the gap to structure-based predictors, and parameter-efficient adaptation adds a further consistent gain. The data efficiency observed on AB-Bind and AbBiBench is the same finding viewed from a different angle. If a pre-trained protein language model has already absorbed the evolutionary statistics that constrain binding, then specialising it to a new antigen should not require relearning interaction biochemistry, only aligning an existing geometry with a specific assay scale. Updating fewer than 1% parameters of the backbone weights on small set of a typical DMS campaign data is enough to recover strong correlation across nine antibody-antigen systems and to surpass methods trained on three times the data on AB-Bind. The fact that this is enough is the strongest piece of evidence that the model is doing the right thing. Residue-level attribution closes the loop between prediction and experiment. Because hotspots are recovered without structural supervision, the same explanation pipeline applies in the regime where structure-based explanation is not available, which is the regime where the predictor is most useful. The maps are a hypothesis-generation tool rather than a docking output. They are local, they cannot by themselves separate direct contacts from allosteric or conformational contributors, and the Pro-27 example in p53 illustrates the kind of residue that the method legitimately flags but that a crystal structure would not place in the contact patch. Used as a cross-check against mutagenesis or structural data, attribution maps reduce reliance on opaque scores and give experimentalists a per-residue rationale to act on before committing assay capacity. The consistent localisation of high-attribution residues to interfacial regions across enzyme-inhibitor, signalling, toxin-immunity and antibody-antigen systems indicates that the model is picking up biophysical determinants of binding such as electrostatic complementarity and hydrophobic anchoring rather than diffuse sequence features. Several limitations bound the present work. Performance still declines under extreme evolutionary divergence, which suggests that the most distant relationships in the proteome will require either additional pre-training data or structural priors that the current sequence-only formulation does not carry. zero-shot transfer to an entirely new assay landscape can exhibit scale mismatch and typically requires calibration on at least small set of assay-specific data, which is small in absolute terms but is not zero. The LoRA rank was selected empirically and was not exhaustively swept, and a systematic study across larger ESM-2 variants and alternative parameter-efficient methods may yield further gains. The Integrated Gradients attributions are computed on mean-pooled embeddings and do not yet provide pairwise residue-residue contact predictions. The benchmark, while larger than most prior sequence-only studies, is still skewed toward well-studied protein families, and prospective deployment on truly orphan scaffolds remains to be tested. The model addresses equilibrium affinity rather than kinetics, and post-translational modifications, non-canonical residues, disordered regions and glycan-mediated interfaces sit outside the current input vocabulary. Two extensions are the natural next steps. The first is active learning on top of the few-shot adaptation, in which model uncertainty selects the most informative variants for the next assay batch and reduces the data requirement further^17,41,42^. For an antibody campaign in which a single SPR run already takes the zero-shot correlation from negative to above 0.7, an uncertainty-driven loop is the route to making the data requirement smaller still. The second is the integration of attention-derived contact maps with the attribution output, to move from per-residue importance toward pairwise interface predictions without giving up the sequence-only nature of the model. By coupling a metric-learning view of binding with parameter- efficient adaptation and per-residue explanation, BALM-PPI provides a sequence-only route to protein-protein affinity prediction that is accurate, transferable and inspectable. To our knowledge, it is the first sequence-only framework to recover experimentally validated interface hotspots across enzyme-inhibitor, signalling, toxin-immunity and antibody-antigen systems without any structural supervision, and to do so within a parameter-efficient adaptation framework that can be specialised to a new target with the data generated by a single binding assay.

## 4 Methods

### 4.1 Dataset

We use the PPB-Affinity dataset ^19^, the largest publicly available collection of curated protein-protein binding affinity data. The dataset has 12062 interactions available out of which we extracted 12019 interaction sequences using PDB IDs, and chain IDs. It is instructive to assert that PPB-Affinity dataset contains data from multiple sources, including SKEMPIv2.0, PDBBind, and SAbDab, covering a wide range of protein families. The dataset statistics are summarized in Supplementary Table S1.

### 4.2 Data splits and cross-validation strategy

To rigorously evaluate the model’s generalization capabilities and mitigate the risk of over-optimistic performance due to data leakage, we implemented three distinct partitioning strategies. Each strategy was evaluated using a 5-fold cross-validation (CV) framework. In this configuration, the dataset was randomly partitioned for *random split* into training (80%) and testing (20%) sets, resulting in approximately 9,615 training and 2,404 test samples per fold. This split measures the model’s ability to interpolate within the known distribution of the dataset, where similar proteins or identical complexes may appear in both train and test sets. Evaluating models using random split often results in data leakage, leading to overestimating the model’s generalizability. ^20^ To simulate a more realistic drug discovery scenario where the model must predict affinities for novel protein complexes, we implemented a *cold split* using the GroupKFold algorithm. By grouping the data according to PDB IDs, we ensured that all instances of a specific protein complex were assigned exclusively to either the training or the testing set. This prevents the model from memorizing specific structural configurations and requires it to generalize to entirely unseen protein-protein interfaces. The most challenging evaluation involves predicting interactions for evolutionarily distant proteins. To achieve this, we employed *sequence-similarity split* where protein sequences are clustered based on homology to eliminate redundancy. We calculated pairwise sequence similarity using *K*-mer Jaccard similarity (*K* = 3) ^43^. An agglomerative clustering algorithm with an average-linkage criterion was applied to the pre-computed distance matrix (*Distance* = 1 − *Similarity*). A strict similarity threshold of 30% was enforced; sequences within a cluster share higher similarity, while separate clusters are evolutionarily divergent^44^. We applied a strict pair-exclusion rule, if either the target or the partner protein in a pair belonged to a test-assigned cluster, the entire interaction was moved to the test set. This strategy ensures that the model cannot rely on homologous sequences seen during training, providing a true measure of its ability to generalize across novel protein families.

### 4.3 Architectural framework for affinity prediction

To rigorously test our hypothesis that a similarity learning objective is superior for affinity prediction, we designed and compared three distinct architectural paradigms. These architectures share the same foundational protein encoder (ESM-2) but differ fundamentally in their approach of integrating information and making predictions. We first implemented a standard regression baseline, which is widely adopted and is a powerful regression strategy. This design is on the idea of learning a complex function from the complete set of features of the interacting pair. It first generates a 1280-dimensional embedding for each protein via ESM-2.The core of this approach lies in its information fusion step under which the both interacting protein embeddings are concatenated into a single, high-dimensional feature vector. This vector, which represents the protein pair as a single entity, is then fed into a Multi-Layer Perceptron (MLP) with linear projection, ReLU, dropout, and output layer. The MLP is trained to directly regress the *pK*_*d*_value. While this method is conceptually simple and effective, its primary limitation is that it learns general features of the pair, rather than creating reusable, independent representations for each protein.

To better respect the inherent symmetry of protein–protein interactions, we developed BALM-PPI. This framework reframes affinity prediction as a metric learning problem. We hypothesized that the symmetric nature of protein-protein binding could be more effectively modeled by learning a shared binding space where alignment directly correlates with affinity. We pass two protein sequences to ESM-2 encoding, and each embedding is transformed by its own Projection Layer into a 256-dimensional latent space. Crucially, these projected vectors are then L2-normalized. The final prediction is then determined by the cosine similarity between these two normalized vectors. To fully realize the potential of our model’s framework, our final and best-performing implementation, BALM-PPI, as illustrated in Fig. 1, integrates PEFT with our proposed framework. By injecting trainable LoRA adapters into the ESM-2 backbone, the language model itself learns to produce embeddings that are optimized for this specific binding affinity task, creating a powerful synergy between the feature extractor and the learning objective.

#### 4.3.0.1 Protein Encoding Layer

Each interaction partner is processed through the ESM-2 transformer ^9^. To accurately model multi-chain proteins (e.g., Antibody Heavy/Light chains or TCR *α*/*β* chains), we implement a sequence-level delimiter strategy. We utilize two consecutive CLS tokens as structural delimiters at chain boundaries. This dual CLS token configuration enables the transformer’s self-attention layers to explicitly recognize structural discontinuities and differentiate between distinct polypeptide chains while maintaining a unified representational pass.^45^ This substitution allows the transformer’s self-attention mechanism to recognize chain boundaries and structural breaks within a single encoding pass.

Following tokenization, the hidden states from the final layer (*L* = 33) are extracted. We apply a masked mean-pooling strategy over the sequence length, ensuring that padding tokens are excluded from the representation. Formally, for a processed sequence *S*_*i*_ with hidden states **h**_*i*, *j*_, embedding **e**_*i*, *j*_ is defined below:

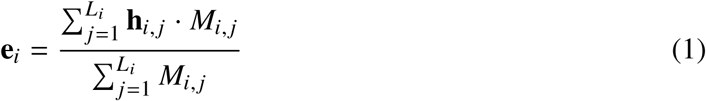

here *M*_*i*, *j*_ is a binary attention mask. This yields a fixed-length 1280-dimensional vector representing the biophysical state of the entire protein complex on one side of the interaction.

#### 4.3.0.2 Projection to Shared Latent Space

We project each protein’s ESM-2 embedding into a 256-dimensional shared latent space using a two-stream architecture. This approach employs separate, independently trained linear layers for each protein, allowing the model to learn potentially distinct transformations.The projected embeddings for protein 1 and protein 2 are calculated using the following expression :

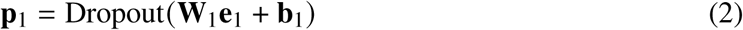

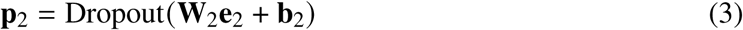

where **W**_1_, **W**_2_ ∈ R^256×1280^ are the projection matrices and the dropout rate is set to 0.1. This design offers greater flexibility than a weight-sharing siamese approach.

#### 4.3.0.3 L2 Normalization and Cosine Similarity

Both projected embeddings p1,p2 from equation 2 and 3 respectively are L2-normalized to unit vectors, enabling stable cosine similarity computation using the following expressions:

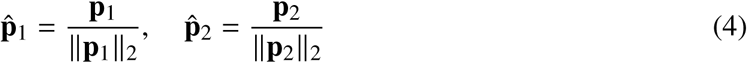

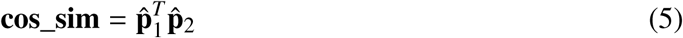

#### 4.3.0.4 *pK*_*d*_ Prediction and Loss Function

Cosine similarity values (cos_sim) which are in the range [-1, 1] are mapped to the dataset’s *pK*_*d*_ range through linear transformation using following equation:

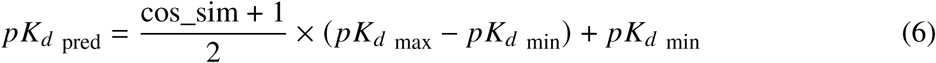

The model is trained using cosine MSE loss, where ground truth *pK*_*d*_ values are scaled to [-1, 1] for cosine similarity learning:

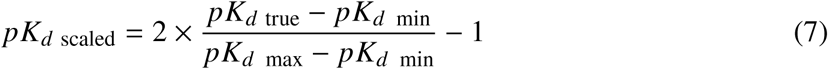

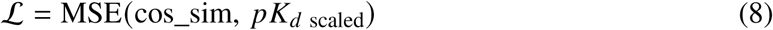

To improve the performance of BALM-PPI without PEFT and make it more robust and efficient, we propose an enhancement over this model. The new architecture named BALM-PPI employs PEFT using LoRA adapters. Instead of fully fine-tuning the ESM-2 model, we train less than 1% of the parameters by injecting these adapters into the layers of ESM-2. This approach updates only the adapter weights while keeping the remaining model architecture unchanged. The results of BALM-PPI are encouraging and are discussed in the result section in details. The following subsections elicits some of the components of the BALM-PPI.

#### 4.3.1 Parameter-Efficient Fine-Tuning (PEFT)

To adapt the ESM-2 model to the nuances of protein binding energetics without full-parameter retraining, we introduce BALM-PPI. This framework integrates PEFT through Low-Rank Adaptation (LoRA) within the metric learning objective.

We inject trainable low-rank update matrices into the frozen transformer backbone, specifically targeting the Query (*Q*), Key (*K*), and Value (*V*) projection modules of the self-attention mechanism. For a weight matrix *W* ∈ R^*d*×*k*^, the update is formulated as:

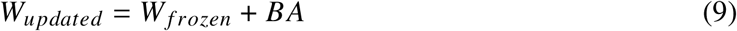

where *B* ∈ R^*d*×*r*^ and *A* ∈ R^*r*×*k*^ are trainable low-rank matrices. In our implementation, we utilize a rank *r* = 8 and a scaling factor *α* = 16. This approach allows BALM-PPI to specialize the language model’s internal attention on binding hotspots by updating only 0.31% (approx. 2.02 million) of the total parameters. Crucially, as shown in Fig. 1, the two-stream architecture allows the fine-tuned model to project these multi-chain aware embeddings into a shared latent space where the cosine similarity is optimized to correlate with experimental *pK*_*d*_ values. PEFT provides a way to provide modular fine-tuning with optimal cost and resources. We integrate Low Rank Adaptation (LoRA) adapters into ESM-2 attention layers to enable parameter-efficient adaptation. LoRA fine-tunes models by injecting small low-rank update matrices into frozen weights, instead of retraining all weights ^15^. With PEFT enabled, less than 1% of model parameters are required to be trained which, dramatically reduces computational requirements while maintaining representation quality.

### 4.4 Quantifying residue contributions to binding affinity

To better understand how individual residues influence predicted binding affinity, we incorporated an explainability framework based on the PyTorch Captum library (Version- 0.7.0) through the Integrated Gradients^18^ method. IG attribute the output prediction to each input feature by integrating the gradients of the model’s output along a straight line from a baseline (zero embedding) to the actual input embedding. This provides a robust measure of how each residue influences the model’s predicted affinity (*pK*_*d*_)).

For each protein pair,two protein sequences *S*_1_ and *S*_2_ were independently tokenized and embedded using the pretrained ESM-2 language model with 650-million-parameters. During attribution computation, one protein was fixed while IG attributions were calculated for the other, and vice versa, yielding residue-level importance scores for both proteins.

Formally, for a given sequence *S* = {*x*_1_, *x*_2_, . . ., *x*_*n*_}, the attribution *A*_*i*_ for residue *i* is computed as:

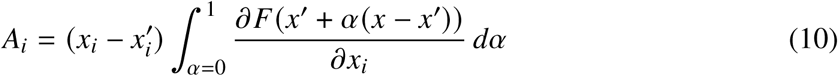

where *x* and *x*^′^ denote the actual and baseline embeddings, respectively, and *F*(·) is the model’s cosine similarity output from the BALM-PPI projection head. All attributions were normalized such that their absolute sum per sequence equals one, enabling comparability across proteins. It is important to note that these attributions represent the *individual importance* of each residue to the overall predicted binding affinity, computed independently for each protein. BALM-PPI uses mean-pooled protein embeddings before computing similarity. Therefore, our attributions quantify how much each residue contributes to its protein’s overall binding propensity.This approach is particularly suited to our metric learning framework, where binding affinity emerges from the relationship between protein-level representations.

## 5 Author contributions

All authors have contributed equally to this work.

## 7 Data availability

All the datasets used for this study are publicly available under :

**PPB-Affinity datase:** https://zenodo.org/doi/10.5281/zenodo.11070823

**AB-Bibench assays:** https://huggingface.co/datasets/AbBibench/Antibody_Binding_Benchmark_Dataset

**AB-Bind:** https://github.com/TAI-Medical-Lab/MVSF-AB

The pretrained model is accessible on Hugging Face at https://huggingface.co/Harshit494/BALM-PPI

## 8 Code availability

**BALM-PPI implementation:** https://github.com/rgorantla04/BALM-PPI.git

## 9 Reporting summary

**Supplementary Information:** All other detailed information provided in supplementary materials.

## 10 Acknowledgment

Harshit Singh would like to acknowledge the support of Department of Computer Science and Engineering, Shiv Nadar University, Delhi-NCR for providing all the resources needed for computation. Rohan Gorantla acknowledges the support and funding of the United Kingdom Research and Innovation (grant EP/S02431X/1), UKRI Centre for Doctoral Training in Biomedical AI at the School of Informatics, University of Edinburgh, and Exscientia Plc, Oxford.

## Supplementary Information

### Contents

This Supplementary Information contains dataset composition analyses, expanded benchmarking results, few-shot adaptation experiments, architecture and backbone ablations, web-server outputs, and additional residue-level explainability case studies. References cited in the Supplementary Information are called directly from the shared references.bib database and listed separately at the end of the Supplementary Information.

#### S1 Supplementary Note 1: Dataset Characteristics and Preprocessing

The PPB-Affinity dataset was compiled from five primary sources: SKEMPI v2.0, SAbDab, Affinity Benchmark v5.5, PDBbind v2020, and ATLAS^1^. To provide a comprehensive overview of the PPB-Affinity dataset, we analyzed the distribution of protein structures and experimental affinity values. The dataset is derived from five major repositories, each exhibiting distinct affinity profiles (Fig. S1b). The distribution of the top 20 PDB structures (Fig. S1a) reveals that while certain well-studied complexes (e.g., 1A07, 3SGB) are highly represented, the dataset maintains a broad structural diversity. The overall distribution of binding affinity (*pK*_*d*_) values follows a Gaussian-like distribution centered at approximately 7.55 (Fig. S1c). For the purposes of downstream analysis and explainability, we categorized the interactions into Binders (*pK*_*d*_ ≥ 7) and Non-binders (*pK*_*d*_ < 7). The density plots in Fig. S1d demonstrate clear separation between these classes, while maintaining sufficient overlap to challenge the model’s regression capabilities.

**Figure S1.**
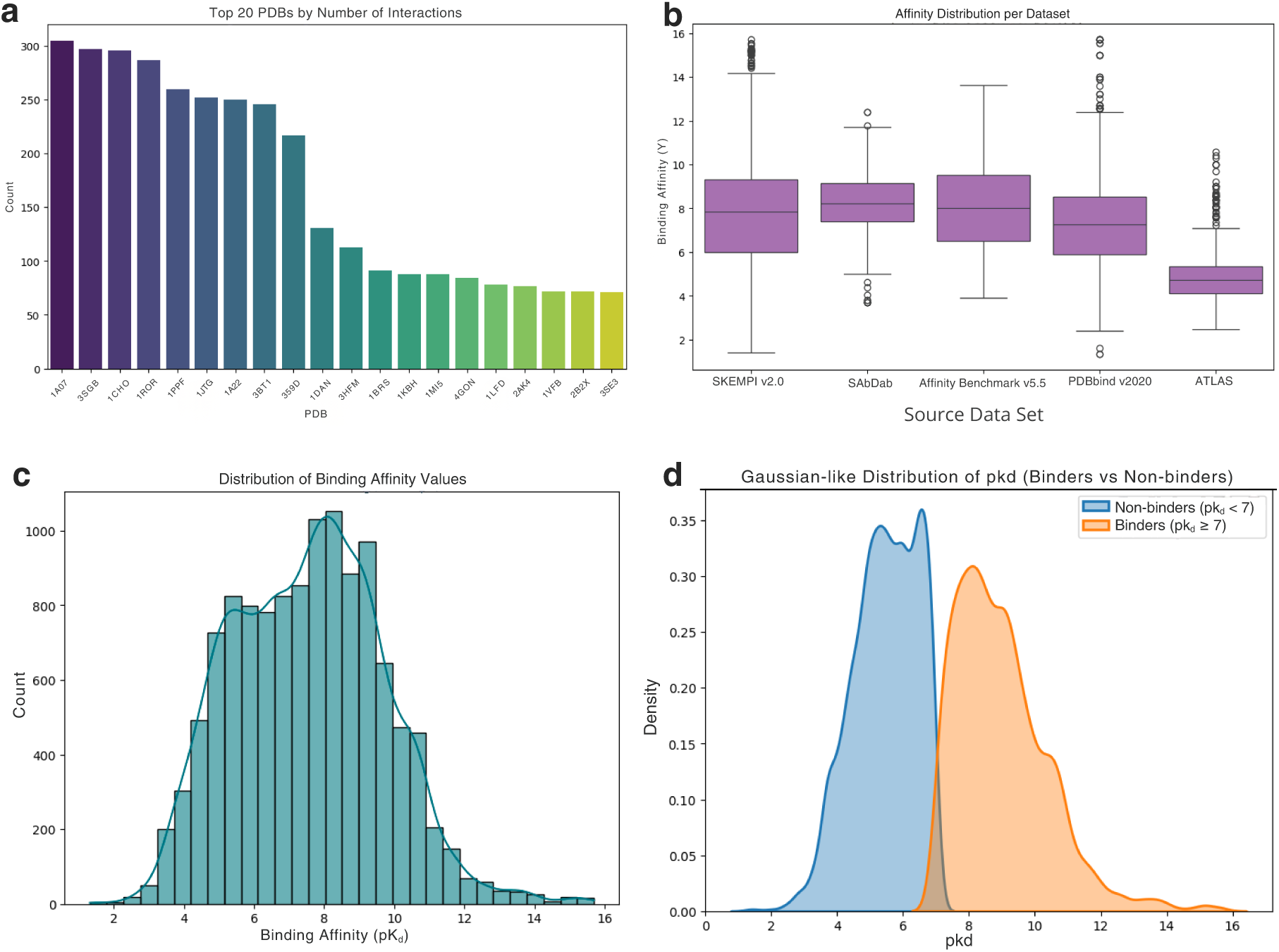
Overview of the PPB-Affinity dataset composition. **a**, Top 20 PDB identifiers by number of interactions, illustrating the representation of common structural scaffolds. **b**, Distribution of experimental binding affinities (*pK*_*d*_) across the five source datasets, showing median values and outliers. **c**, Global distribution of binding affinity values across the entire processed dataset (*n* = 12, 019). **d**, Density distribution of binding affinities split by binder status, showing a Gaussian-like profile for both populations used in model calibration.

Table S1 provides the breakdown of the 12,019 interactions used in this study after filtering for sequence availability and quality.

**Table S1.** PPB-Affinity dataset statistics after preprocessing.

| Attribute | Value |
| --- | --- |
| Total interactions | 12,019 |
| Unique PDB structures | 3,030 |
| Antibody-antigen complexes | 1,395 |
| TCR-pMHC complexes | 1,431 |
| Other protein complexes | 9,193 |
| $pK_d$ range | 1.32 – 15.70 |
| $pK_d$ mean ± s.d. | 7.55 ± 2.15 |

#### S2 Supplementary Note 2: Expanded Benchmarking Performance

To evaluate the robustness of BALM-PPI, we compared it against a standard regression baseline and a version of the model without Parameter-Efficient Fine-Tuning (PEFT).^2^ Table S2 details the performance across three validation splits. Furthermore, we compared our sequence-only approach against the structure-based PPB-Affinity baseline (Table S3) and analyzed performance across different database sources (Table S4).

**Table S2.** Split-wise performance comparison (mean ± error). Lower RMSE and higher Pearson/Spearman indicate better performance.

| Model | Split | RMSE ↓ | Pearson ↑ | Spearman ↑ |
| --- | --- | --- | --- | --- |
| Baseline | Random | 1.345 ± 0.029 | 0.779 ± 0.001 | 0.773 ± 0.014 |
| | Cold | 1.631 ± 0.022 | 0.650 ± 0.022 | 0.634 ± 0.043 |
| | Sequence | 1.755 ± 0.160 | 0.555 ± 0.097 | 0.545 ± 0.103 |
| BALM-PPI without PEFT | Random | 1.192 ± 0.022 | 0.833 ± 0.007 | 0.832 ± 0.007 |
| | Cold | 1.545 ± 0.068 | 0.698 ± 0.023 | 0.691 ± 0.021 |
| | Sequence | 1.717 ± 0.195 | 0.583 ± 0.110 | 0.566 ± 0.115 |
| BALM-PPI | Random | **0.994 ± 0.025** | **0.887 ± 0.008** | **0.886 ± 0.008** |
|  | Cold | **1.486 ± 0.115** | **0.730 ± 0.025** | **0.715 ± 0.046** |
|  | Sequence | **1.672 ± 0.195** | **0.612 ± 0.086** | **0.602 ± 0.087** |

**Table S3.** Performance comparison of BALM-PPI (Sequence) vs. the PPB-Affinity baseline (Structure). Best performance for each metric is highlighted in bold. BALM-PPI results are reported as mean ± error.

| Subgroup | Metric | PPB-Affinity (Structure) | BALM-PPI (Sequence) |
| --- | --- | --- | --- |
| Entire | RMSE ↓ | 2.104 | **1.486 ± 0.115** |
| | Pearson ↑ | 0.70 | **0.730 ± 0.025** |
| | Spearman ↑ | 0.691 | **0.715 ± 0.046** |
| TCR-pMHC | RMSE ↓ | 1.615 | **1.045 ± 0.077** |
| | Pearson ↑ | 0.443 | **0.586 ± 0.039** |
| | Spearman ↑ | **0.452** | 0.380 ± 0.049 |
| Antigen-Antibody | RMSE ↓ | 1.994 | **1.286 ± 0.024** |
| | Pearson ↑ | 0.329 | **0.490 ± 0.013** |
| | Spearman ↑ | 0.298 | **0.486 ± 0.032** |

**Table S4.** Source-wise performance comparison of BALM-PPI (Sequence) vs. PPB-Affinity (Structure) across different data sources. BALM-PPI results are reported as mean ± std.

| Source | BALM-PPI (Sequence) |  |  | PPB-Affinity (Structure) |  |  |
| --- | --- | --- | --- | --- | --- | --- |
| | RMSE ↓ | Pearson ↑ | Spearman ↑ | RMSE ↓ | Pearson ↑ | Spearman ↑ |
| SKEMPI v2.0 | 1.549 ± 0.087 | 0.745 ± 0.022 | 0.719 ± 0.022 | 2.103 | 0.744 | 0.741 |
| SAbDab | 1.282 ± 0.033 | 0.490 ± 0.020 | 0.475 ± 0.035 | 1.967 | 0.355 | 0.327 |
| Affinity Benchmark v5.5 | 1.482 ± 0.087 | 0.702 ± 0.031 | 0.692 ± 0.043 | 2.207 | 0.671 | 0.658 |
| PDBbind v2020 | 1.466 ± 0.028 | 0.664 ± 0.006 | 0.661 ± 0.008 | 2.216 | 0.662 | 0.660 |
| ATLAS | 0.969 ± 0.109 | 0.617 ± 0.111 | 0.315 ± 0.086 | 1.594 | 0.387 | 0.433 |

#### S3 Supplementary Note 3: Few-Shot Adaptation on Mutational Landscapes

To test the model’s ability to capture fine-grained mutational effects, we utilized nine assays from the AbBiBench benchmark^3^. Table S5 provides details on the antibodies and antigens involved. We observed that while zero-shot performance varies, few-shot adaptation with as little as 10% of data dramatically improves correlation (Table S6, and Figure S2).

**Table S5.**
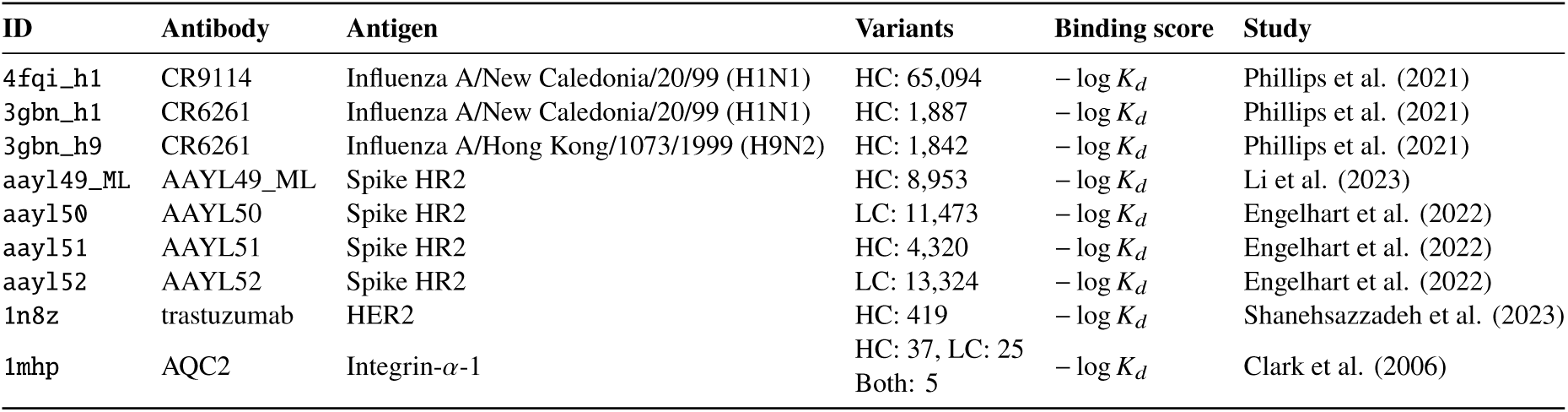
Overview of the binding-affinity assays selected from the AbBiBench benchmark for zero-shot and few-shot evaluation.

**Table S6.**
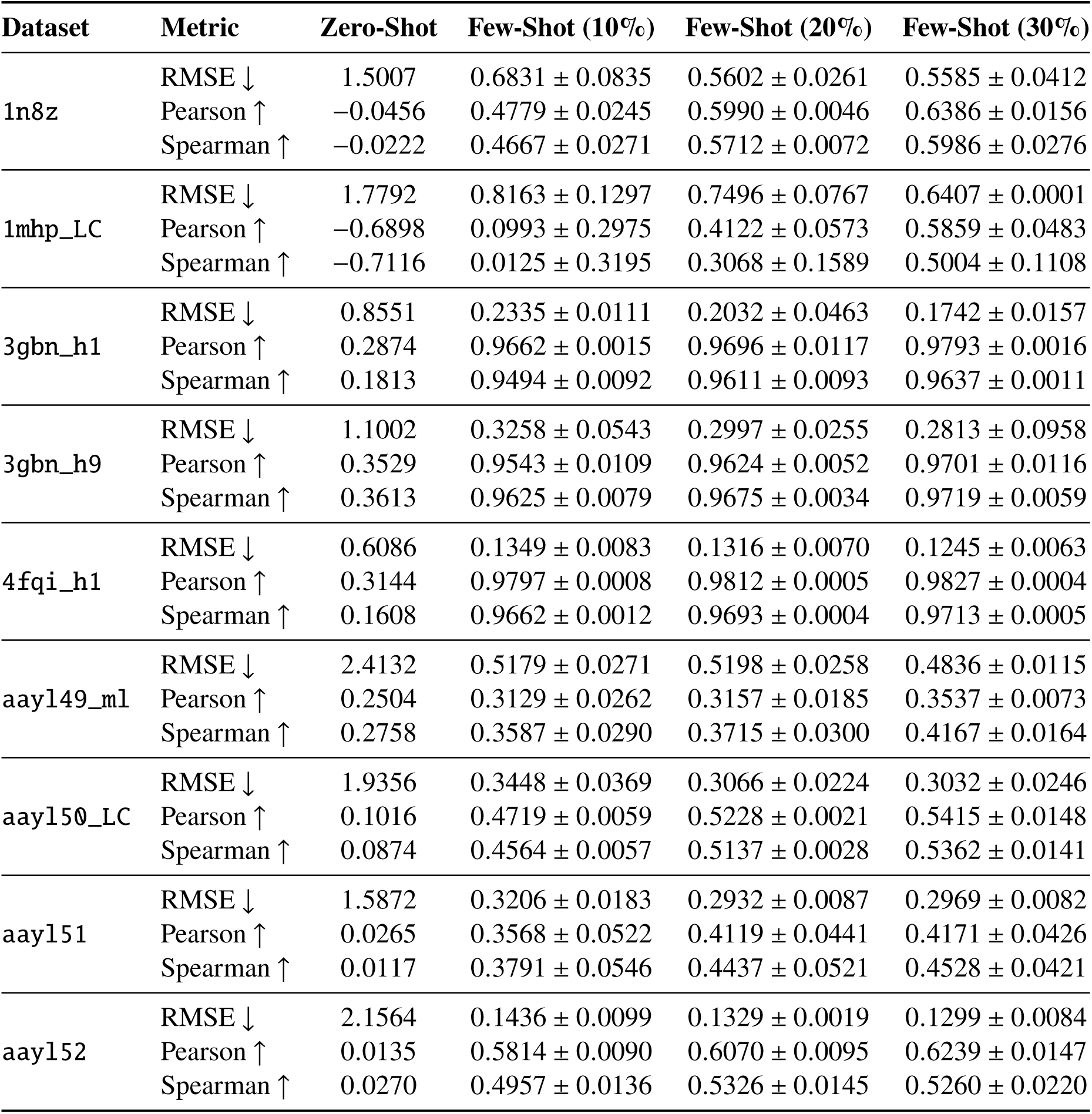
Zero-shot and few-shot performance of BALM-PPI on diverse assays from the AbBiBench benchmark cross diverse datasets. The results demonstrate significant performance gains with minimal task-specific fine-tuning across Influenza, SARS-CoV-2, HER2, and Integrin systems.

| Dataset | Metric | Zero-Shot | Few-Shot (10%) | Few-Shot (20%) | Few-Shot (30%) |
| --- | --- | --- | --- | --- | --- |
| 1n8z | RMSE ↓ | 1.5007 | 0.6831 ± 0.0835 | 0.5602 ± 0.0261 | 0.5585 ± 0.0412 |
|  | Pearson ↑ | −0.0456 | 0.4779 ± 0.0245 | 0.5990 ± 0.0046 | 0.6386 ± 0.0156 |
|  | Spearman ↑ | −0.0222 | 0.4667 ± 0.0271 | 0.5712 ± 0.0072 | 0.5986 ± 0.0276 |
| 1mhp_LC | RMSE ↓ | 1.7792 | 0.8163 ± 0.1297 | 0.7496 ± 0.0767 | 0.6407 ± 0.0001 |
|  | Pearson ↑ | −0.6898 | 0.0993 ± 0.2975 | 0.4122 ± 0.0573 | 0.5859 ± 0.0483 |
|  | Spearman ↑ | −0.7116 | 0.0125 ± 0.3195 | 0.3068 ± 0.1589 | 0.5004 ± 0.1108 |
| 3gbn_h1 | RMSE ↓ | 0.8551 | 0.2335 ± 0.0111 | 0.2032 ± 0.0463 | 0.1742 ± 0.0157 |
|  | Pearson ↑ | 0.2874 | 0.9662 ± 0.0015 | 0.9696 ± 0.0117 | 0.9793 ± 0.0016 |
|  | Spearman ↑ | 0.1813 | 0.9494 ± 0.0092 | 0.9611 ± 0.0093 | 0.9637 ± 0.0011 |
| 3gbn_h9 | RMSE ↓ | 1.1002 | 0.3258 ± 0.0543 | 0.2997 ± 0.0255 | 0.2813 ± 0.0958 |
|  | Pearson ↑ | 0.3529 | 0.9543 ± 0.0109 | 0.9624 ± 0.0052 | 0.9701 ± 0.0116 |
|  | Spearman ↑ | 0.3613 | 0.9625 ± 0.0079 | 0.9675 ± 0.0034 | 0.9719 ± 0.0059 |
| 4fqi_h1 | RMSE ↓ | 0.6086 | 0.1349 ± 0.0083 | 0.1316 ± 0.0070 | 0.1245 ± 0.0063 |
|  | Pearson ↑ | 0.3144 | 0.9797 ± 0.0008 | 0.9812 ± 0.0005 | 0.9827 ± 0.0004 |
|  | Spearman ↑ | 0.1608 | 0.9662 ± 0.0012 | 0.9693 ± 0.0004 | 0.9713 ± 0.0005 |
| aayl49_ml | RMSE ↓ | 2.4132 | 0.5179 ± 0.0271 | 0.5198 ± 0.0258 | 0.4836 ± 0.0115 |
|  | Pearson ↑ | 0.2504 | 0.3129 ± 0.0262 | 0.3157 ± 0.0185 | 0.3537 ± 0.0073 |
|  | Spearman ↑ | 0.2758 | 0.3587 ± 0.0290 | 0.3715 ± 0.0300 | 0.4167 ± 0.0164 |
| aayl50_LC | RMSE ↓ | 1.9356 | 0.3448 ± 0.0369 | 0.3066 ± 0.0224 | 0.3032 ± 0.0246 |
|  | Pearson ↑ | 0.1016 | 0.4719 ± 0.0059 | 0.5228 ± 0.0021 | 0.5415 ± 0.0148 |
|  | Spearman ↑ | 0.0874 | 0.4564 ± 0.0057 | 0.5137 ± 0.0028 | 0.5362 ± 0.0141 |
| aayl51 | RMSE ↓ | 1.5872 | 0.3206 ± 0.0183 | 0.2932 ± 0.0087 | 0.2969 ± 0.0082 |
|  | Pearson ↑ | 0.0265 | 0.3568 ± 0.0522 | 0.4119 ± 0.0441 | 0.4171 ± 0.0426 |
|  | Spearman ↑ | 0.0117 | 0.3791 ± 0.0546 | 0.4437 ± 0.0521 | 0.4528 ± 0.0421 |
| aayl52 | RMSE ↓ | 2.1564 | 0.1436 ± 0.0099 | 0.1329 ± 0.0019 | 0.1299 ± 0.0084 |
|  | Pearson ↑ | 0.0135 | 0.5814 ± 0.0090 | 0.6070 ± 0.0095 | 0.6239 ± 0.0147 |
|  | Spearman ↑ | 0.0270 | 0.4957 ± 0.0136 | 0.5326 ± 0.0145 | 0.5260 ± 0.0220 |

**Figure S2.**
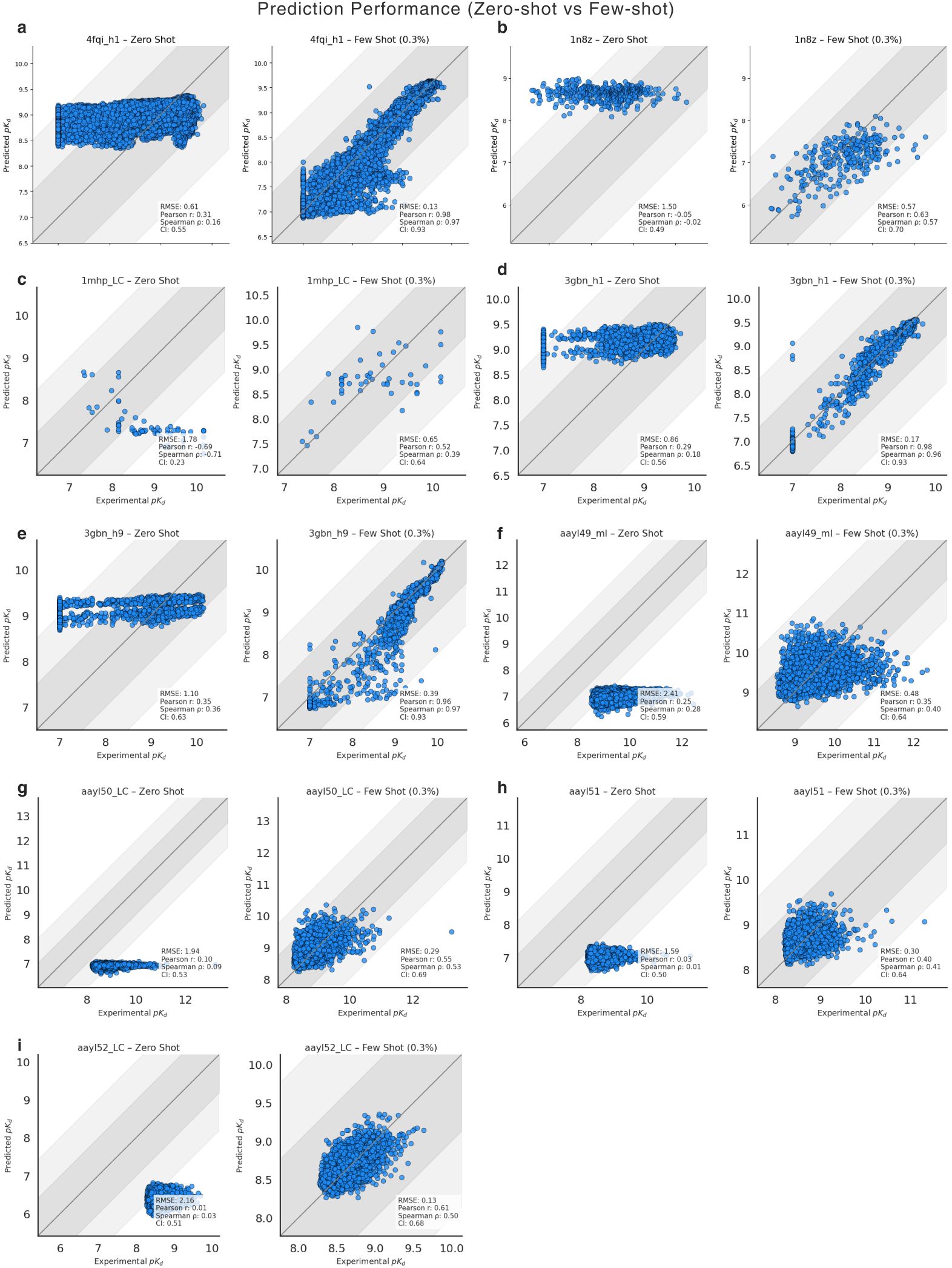
Zero-shot and few-shot performance across AbBiBench mutational landscapes. Scatter plots compare experimental binding scores with BALM-PPI predictions for the nine AbBiBench assays evaluated in Supplementary Note 3. Each panel corresponds to one antibody–antigen assay and contrasts the zero-shot model with few-shot models fine-tuned using 10%, 20%, or 30% of assay-specific data. Points represent individual variants, and tighter alignment with the diagonal indicates improved agreement between predicted and measured binding affinity after few-shot adaptation.

#### S4 Supplementary Note 4: Architecture and Backbone Ablations

We conducted an extensive ablation study to justify the selection of the ESM-2 backbone, comparing ESM-2, ProGen-2, AbLang-2 and ESM-C representations under the same evaluation protocol^4–7^. Table S7 and Fig. S3 illustrates that ESM-2 variants consistently outperform ProGen and AbLang-2 across different projection head sizes.

**Figure S3.**
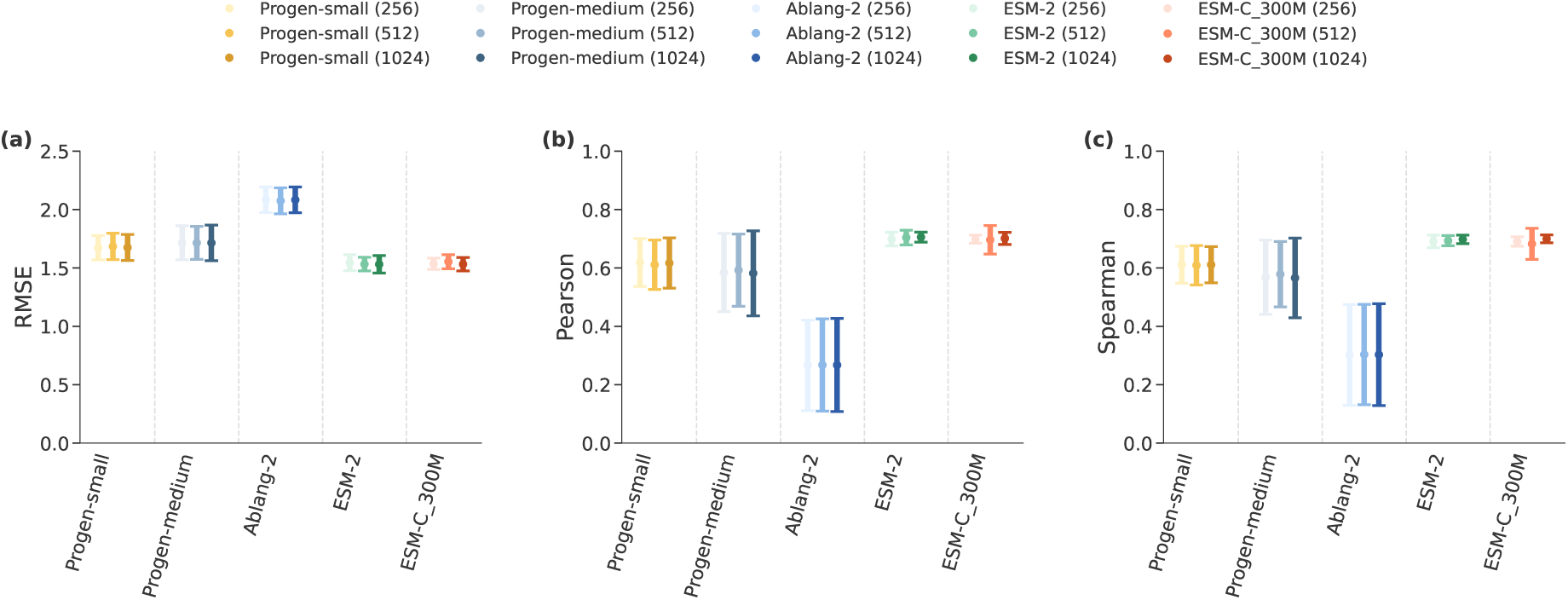
Ablation of protein language model backbones and projection-head capacity. Performance on the cold split for BALM-PPI-style models built on different PLM encoders (ProGen-2 small & medium, AbLang-2, ESM-2, and ESM-C) and different projection-head sizes (256, 512, and 1024). The plot compares backbones under a matched evaluation protocol to isolate the effect of the encoder representation and the capacity of the learned projection into the shared affinity space. The results show that ESM-2 provides the most reliable gains across projection sizes, motivating its use as the default backbone in the main experiments.Error bars show mean ± s.d. across 5 folds.

**Table S7.** Performance across various protein language models and projection sizes under cold split (mean ± error).

| Model | Projection Size | RMSE ↓ | Pearson ↑ | Spearman ↑ |
| --- | --- | --- | --- | --- |
| Progen-small | 256 | 1.673 ± 0.104 | 0.619 ± 0.082 | 0.611 ± 0.064 |
| Progen-small | 512 | 1.684 ± 0.113 | 0.611 ± 0.085 | 0.609 ± 0.068 |
| Progen-small | 1024 | 1.675 ± 0.112 | 0.617 ± 0.086 | 0.611 ± 0.062 |
| Progen-medium | 256 | 1.715 ± 0.146 | 0.584 ± 0.134 | 0.568 ± 0.127 |
| Progen-medium | 512 | 1.715 ± 0.142 | 0.592 ± 0.124 | 0.579 ± 0.112 |
| Progen-medium | 1024 | 1.715 ± 0.152 | 0.582 ± 0.146 | 0.566 ± 0.136 |
| Ablang-2 | 256 | 2.084 ± 0.110 | 0.266 ± 0.155 | 0.302 ± 0.173 |
| Ablang-2 | 512 | 2.075 ± 0.111 | 0.267 ± 0.158 | 0.303 ± 0.172 |
| Ablang-2 | 1024 | 2.083 ± 0.110 | 0.267 ± 0.159 | 0.303 ± 0.174 |
| ESM-2 | 256 | 1.545 ± 0.068 | 0.699 ± 0.024 | 0.691 ± 0.021 |
| ESM-2 | 512 | 1.532 ± 0.059 | 0.704 ± 0.025 | 0.693 ± 0.017 |
| ESM-2 | 1024 | 1.531 ± 0.074 | 0.706 ± 0.017 | 0.698 ± 0.015 |
| ESM-C-300M | 256 | 1.535 ± 0.049 | 0.699 ± 0.014 | 0.690 ± 0.016 |
| ESM-C-300M | 512 | 1.553 ± 0.060 | 0.696 ± 0.049 | 0.682 ± 0.053 |
| ESM-C-300M | 1024 | 1.532 ± 0.057 | 0.701 ± 0.021 | 0.700 ± 0.013 |

#### S5 Supplementary Note 5: BALM-PPI Predict webtool

Supplementary Fig. S4 documents the user-facing BALM-PPI web server that accompanies the model. The interface is organized around two prediction modes: a general protein–protein interaction workflow and an antibody–antigen workflow. In both modes, users can either paste raw amino-acid sequences or retrieve sequences from a PDB identifier and chain selection. The server returns the predicted affinity score, the underlying cosine-similarity value used by the metric-learning model, and an interaction-strength summary calibrated to the training-set *pK*_*d*_scale. When the Integrated Gradients option is enabled, the same prediction run produces residue-level attribution outputs that can be inspected interactively in the browser or downloaded for downstream analysis.

The purpose of Fig. S4 is to show that the explainability component is not a post hoc offline analysis but is integrated directly into the deployment workflow. The NGL viewer maps attribution intensities onto available structural coordinates, while the heatmap and ranked-residue panels provide sequence-level summaries for cases where structures are unavailable or incomplete. The batch-prediction module further supports high-throughput screening by accepting CSV files with multiple sequence pairs and returning downloadable prediction tables, attribution-score CSV files, and JSON summaries. Together, these panels demonstrate how BALM-PPI can be used both for individual mechanistic inspection and for larger candidate-prioritization campaigns.

**Figure S4.**
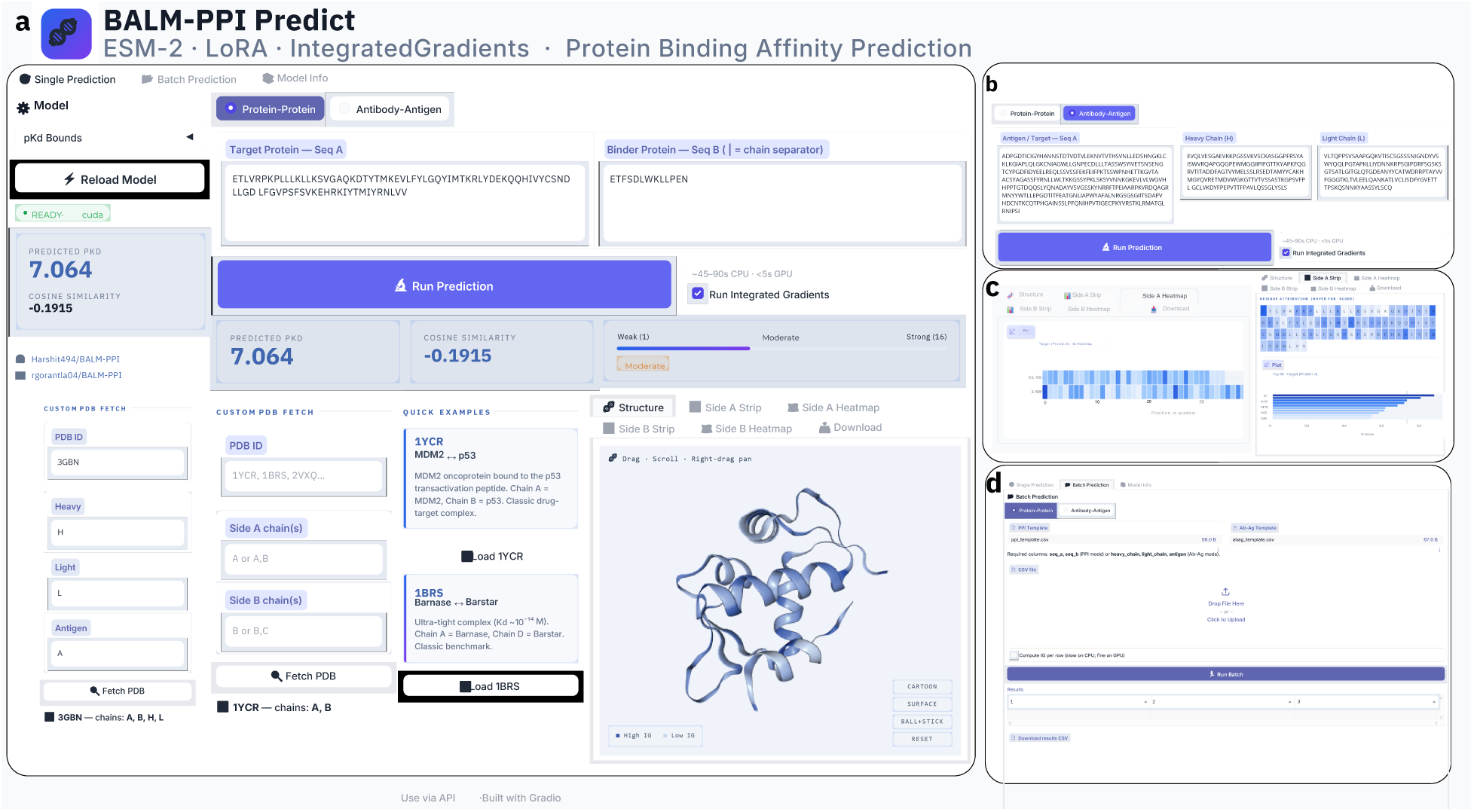
BALM-PPI Predict web server for protein–protein and antibody–antigen binding affinity prediction. **(a)** Protein–protein interaction (PPI) prediction interface. Users may provide target (Seq A) and binder (Seq B) sequences directly or retrieve them from a Protein Data Bank (PDB) structure using PDB and chain identifiers. The server predicts binding affinity (pKd), cosine similarity, and interaction strength, and supports optional Integrated Gradients (IG) analysis for residue-level explainability. An interactive NGL viewer visualizes protein structures with IG-derived residue attributions using cartoon, surface, and ball-and-stick representations. **(b)** Antibody–antigen prediction interface, allowing manual input or PDB-based retrieval of heavy-chain, light-chain, and antigen sequences for affinity prediction and explainability analysis. **(c)** Explainability outputs generated from IG analysis, including residue-level attribution heatmaps for target and binder proteins and ranked plots of the top contributing residues. Attribution scores can be exported as CSV files together with prediction summaries in JSON format. **(d)** Batch prediction module for high-throughput screening. Users can upload template CSV files containing protein–protein or antibody–antigen sequence pairs and perform large-scale affinity prediction, with downloadable results for downstream analysis.

#### S6 Supplementary Note 6: Additional Explainability Case Studies

Supplementary Note 6 expands the residue-level interpretation experiments beyond the condensed main-text figures. The goal is to test whether BALM-PPI highlights physically plausible binding determinants across interaction types that differ in geometry, recognition mechanism, and evolutionary context. For each complex, Integrated Gradients attributions were computed separately for the two interaction partners and visualized both as residue heatmaps and as structural overlays when coordinates were available. Higher-intensity residues indicate positions with larger absolute contribution to the predicted affinity score, rather than experimentally measured energetic changes for single mutations.

Supplementary Fig. S5 focuses on non-antibody protein–protein complexes, including Ras–Raf and BPTI–Trypsin. These systems provide complementary tests of the attribution framework: Ras–Raf represents an extended signalling interface, whereas BPTI–Trypsin is dominated by a compact protease-inhibitor binding geometry. In both cases, the highlighted residues localize to the known interaction surface or to residues supporting the binding pocket, indicating that the sequence-only model assigns importance to mechanistically relevant regions rather than distributing attribution uniformly across the protein sequence.

**Figure S5.**
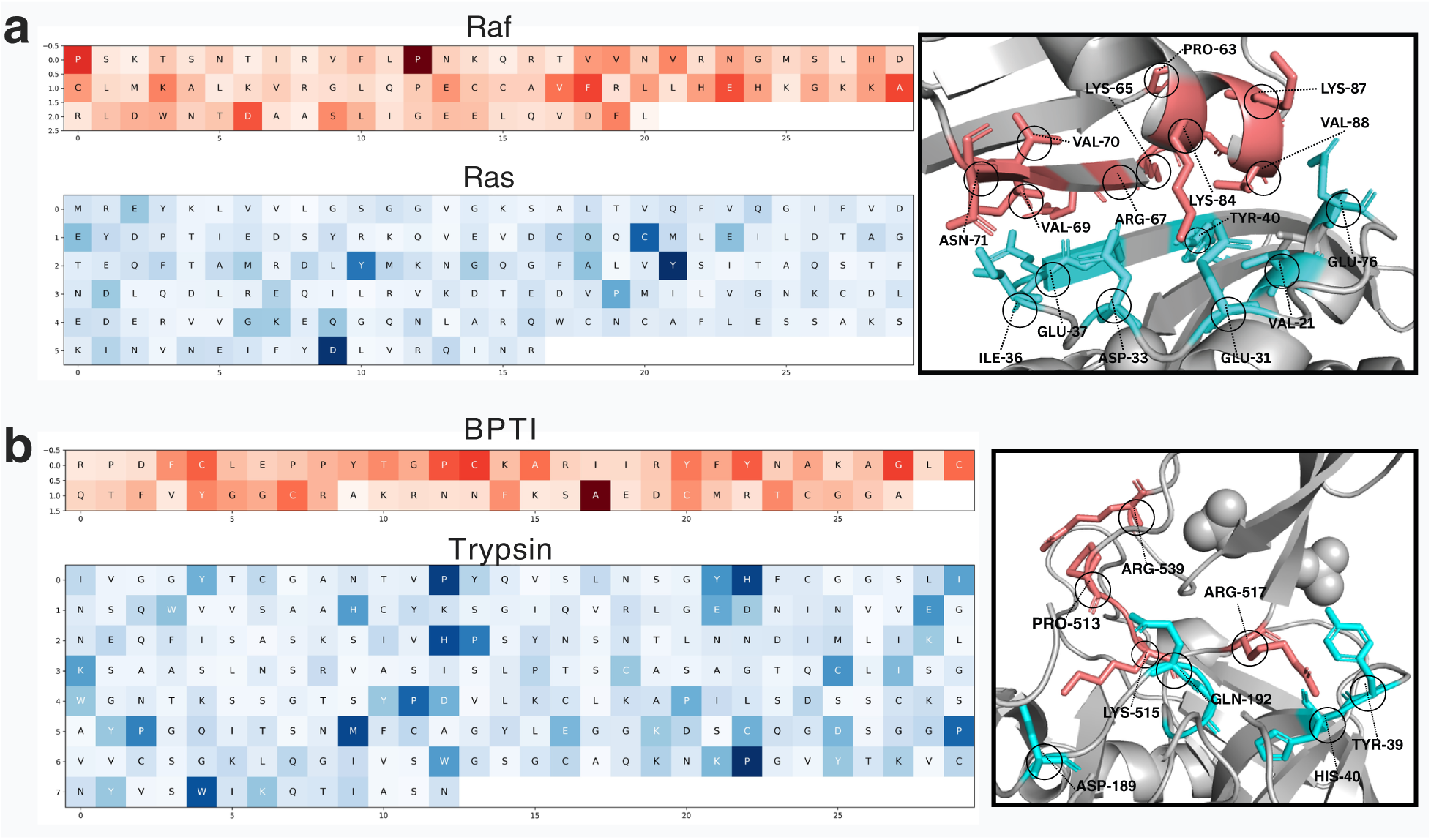
Additional protein-protein interaction explainability examples. Residue-level Integrated Gradients attribution maps and structural visualizations for Ras-Raf and BPTI-Trypsin complexes. These supplementary panels extend the main-text explainability analysis by showing interaction systems that are discussed in the manuscript but displayed in the SI for readability. Higher-intensity residues indicate larger attribution magnitude and stronger contribution to the BALM-PPI affinity prediction.

Supplementary Fig. S6 extends the antibody–antigen analysis to additional influenza and lysozyme recognition examples. These cases evaluate whether BALM-PPI can recover paratope and epitope regions in systems with distinct antibody binding modes, including HA-stem recognition, HA-head recognition, and a canonical HyHEL-10–lysozyme interface. The supplementary panels show that zero-shot attributions generally identify broad interface regions, while the few-shot setting sharpens the attribution signal around assay-relevant residues. These examples support the use of BALM-PPI attributions as a practical guide for variant prioritization and structural cross-checking before experimental follow-up.

**Figure S6.**
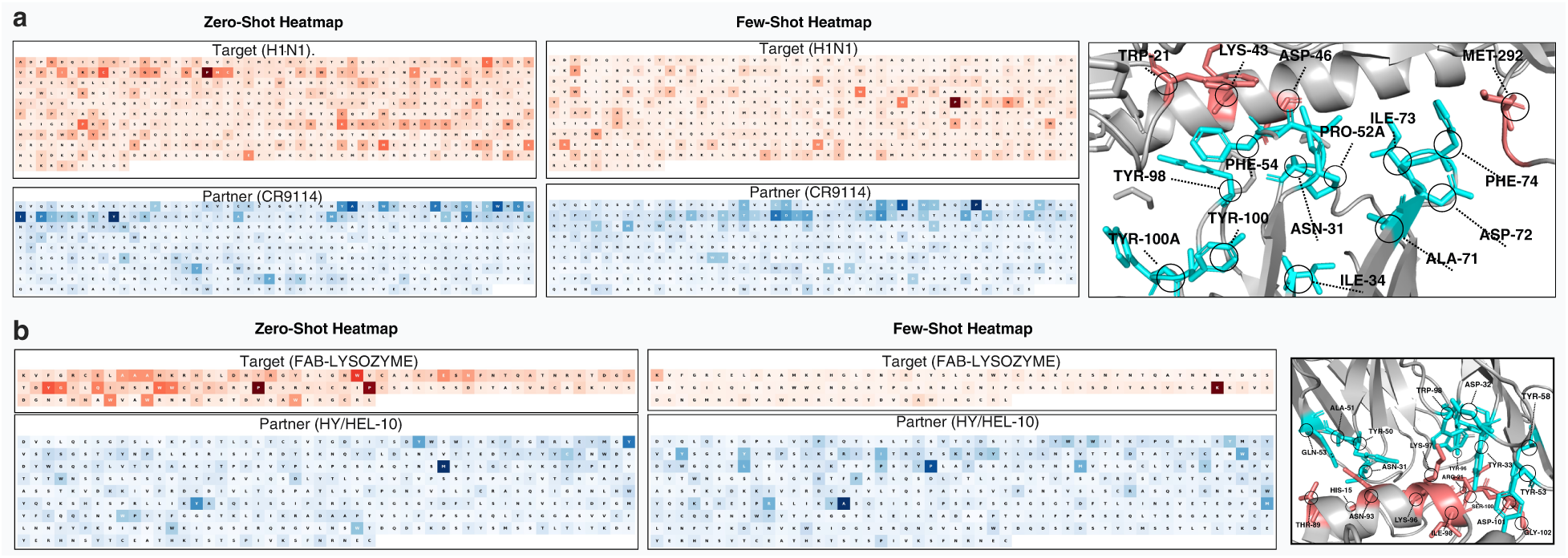
Additional antibody-antigen explainability examples. Residue-level Integrated Gradients heatmaps and structural visualizations for H1N1-related antibody-antigen systems and the HyHEL-10–lysozyme complex, including Target H1N1 with Partner CR9114, Partner HyHEL-10, and Target Fab/Lysozyme panels. These supplementary panels extend the main antibody-antigen explainability figure after CR9114-H1N1, Fab-2D1-H1N1, and HyHEL-10–lysozyme examples were moved from the main figure. Higher-intensity residues indicate larger attribution magnitude and stronger contribution to the BALM-PPI affinity prediction.

## Supplementary References

